# Nanog safeguards early embryogenesis against global activation of maternal β-catenin activity by interfering with TCF factors

**DOI:** 10.1101/821702

**Authors:** Mudan He, Ru Zhang, Fenghua Zhang, Shengbo Jiao, Ding Ye, Houpeng Wang, Yonghua Sun

## Abstract

Maternal β-catenin activity is essential and critical for dorsal induction and its dorsal activation has been thoroughly studied. However, how the maternal β-catenin activity is suppressed in the non-dorsal cells remains poorly understood. Nanog is known to play a central role for maintenance of the pluripotency and maternal-to-zygotic transition. Here we reveal a novel role of Nanog as a strong repressor of maternal Wnt/β-catenin signaling to safeguard the embryo against hyper-activation of maternal β-catenin activity and hyper-dorsalization. Knockdown of *nanog* at different levels led to either posteriorization or dorsalization, mimicking zygotic or maternal activation of Wnt/β-catenin activities, and the maternal-zygotic mutant of *nanog* (MZ*nanog*) showed strong activation of maternal β-catenin and hyper-dorsalization. Although a constitutive-activator-type Nanog (Vp16-Nanog, lacking the N-terminal) perfectly rescued the defects of maternal to zygotic transition in MZ*nanog*, it did not rescue the phenotypes resulting from β-catenin activation. Mechanistically, the N-terminal of Nanog directly interacts with TCF and interferes with the binding of β-catenin to TCF, thereby attenuating the transcriptional activity of β-catenin. Therefore, our study establishes a novel role for Nanog in repressing maternal β-catenin activity and demonstrates a transcriptional switch between β-catenin/TCF and Nanog/TCF complexes, which safeguards the embryo from global activation of maternal β-catenin activity.

## Introduction

The Wnt/β-catenin signaling pathway, known as the canonical Wnt signaling pathway, is highly conserved during evolution. It plays crucial roles in organogenesis, stem cell renewal and differentiation, homeostasis, reproduction, carcinogenesis and embryonic development [1–5]. Decades of studies have shown that the central scheme of the Wnt/β-catenin pathway is to stabilize the transcription coactivator β-catenin and protect it from phosphorylation-dependent degradation [6, 7]. The Wnt/β-catenin pathway is well-known for its ON/OFF regulation model. In the absence of Wnt ligand, cytoplasmic β-catenin forms a complex with Axin, APC, GSK3 and CK1, and is phosphorylated by CK1 and GSK3, recognized by the E3 ubiquitin ligase subunit β-Trcp, and processed through ubiquitination and proteasomal degradation. At this point, the nuclear TCFs is believe to physically interact with the co-repressors (the Groucho/TLE family members) and act as transcriptional repressors [8–11]. In the present of Wnt ligand, a receptor complex forms between Frizzled and LRP5/6, and Frizzled is recruited by Dvl which leads to LRP5/6 phosphorylation and Axin recruitment, which in turn disrupts Axin-mediated phosphorylation/degradation of β-catenin, allowing β-catenin to accumulate in the nucleus where it serves as a coactivator for TCF to activate Wnt-responsive genes [12, 13]. Thus, the activity of Wnt/β-catenin pathway is considered to be directly related to the level of nucleus β-catenin and its interaction with co-activators (such as TCFs) as well as co-repressors (Groucho/TLE) [12, 14].

The dorsal accumulation of maternal β-catenin proteins plays a pivotal role in the dorsal axis formation, and the factors and mechanisms controlling maternal nuclear β-catenin accumulation and activation have been thoroughly studied. It is generally believed that the maternally inherited β-catenin proteins are translocated from the vegetal pole to the future dorsal side of embryos through a microtubule-dependent manner [15, 16]. It was reported that the nuclear β-catenin is stabilized and activated by Wnt ligands, such as Wnt11 in Xenopus [17] and Wnt8a in zebrafish [18]. However, a recent *wnt8a* knockout study in zebrafish suggests that the maternally deposited *wnt8a* mRNA is not required for maternal β-catenin activation [19]. Several studies also indicate that the Wnt receptors are not essentially required for maternal Wnt/β-catenin signaling activation, since knockdown or knockout of Wnt receptor LRP5 did not lead to any early developmental abnormality in mice and zebrafish [20, 21], and dorsal overexpression of dominant-negative LRP6 did not perturb the axis formation in *Xenopus* [22]. A recent knockout study shows that the maternal clearance of Dvl activities did not lead to any early dorsal defects in zebrafish [23], indicating that the maternal β-catenin activation is independent of Wnt ligand-receptor mediated process. More recently, it is demonstrated that a novel membrane protein Huluwa (Hwa) interacts with Axin to interfere the formation of β-catenin destruction complex, thus promoting the activity of maternal β-catenin and dorsal axis formation [24].

In principle, the maternal β-catenin activity should be strictly restricted to the dorsal-most cells and be globally repressed in the embryonic cells other than the dorsal-most cells, since ectopic activation of the maternal β-catenin activity would lead to disrupted dorso-ventral axis. In contrast to numerous studies focusing on the proper activation of maternal β-catenin signaling, the dorsal restriction of β-catenin signals and its repression in non-dorsal cells are poorly understood. For instance, although *axin1* is maternally expressed in zebrafish, the *axin1* mutant, *materblind* (*mbl*) only exhibits a zygotic Wnt/β-catenin activation phenotype in which eyes and telencephalon are reduced or absent and diencephalic fates expand to the front of the brain [25]. A Wnt/β-catenin pathway antagonist Chibby has shown to physically interact with β-catenin to repress WNT signaling [26, 27], whereas, loss of Chibby only results in lung development defect in mice and basal body formation and ciliogenesis defects in *Drosophila*. Another type of β-catenin repressor, *ctnnbip1* (inhibitor of β-catenin and TCF, previously named ICAT) could bind to the C-terminal region of β-catenin to disturb the interaction between β-catenin and TCF, and injection of dominant-negative *ctnnbip1* in ventral side can induce secondary axis in *Xenopus* [28], suggesting that ventral repression of β-catenin activity is crucial for proper dorso-ventral axis formation. However, no mutant analysis of *ctnnbip1* has been described. The protein lysine demethylase Kdm2a/b is reported to mediate the demethylation of β-catenin and to promote its ubiquitination and degradation, but knockdown of *kdm2a/b* only leads to loss of posterior structures in *Xenopus* [29], and *kdm2* homozygous mutants show no developmental defects in *Drosophila* [30, 31]. In zebrafish, Lzts2, Amotl2 and Tob1 are reported to inhibit β-catenin transcriptional activity by physically associating with β-catenin and preventing the formation of β-catenin/LEF1 complexes, but no mutant phenotypes have been reported to show that how these factors affect the dorsal axis formation [32–34]. Therefore, what factor and which kind of mechanism repress the global β-catenin activation in nucleus need to be further studied with genetic null mutants.

Nanog is a core factor for maintenance of pluripotency and self-renewal of embryonic stem (ES) cells [35, 36]. Nanog was first identified in mouse ES cells, and the *nanog*-deficiency ES cells loss pluripotency and *nanog* mutant mice showed defects on ectodermal development and inner cell mass (ICM) proliferation [37]. Although Nanog is not on the list of the classical Yamanaka factors for induced pluripotent stem (iPS) cells, it has been demonstrated to be a “master switch” in the acquisition of cell pluripotency [38]. In zebrafish, by using morpholino-mediated knockdown approaches, previous studies have shown that *nanog* is a pivotal maternal factor to mediate endoderm formation through the Mxtx2-Nodal signaling [39] and initiates the transcription of the first wave of zygotic genes and induces the clearance of maternal mRNAs together with Pou5f3 and SoxB1 during maternal-zygotic transition (MZT) [40]. Recently, by generating maternal zygotic mutants of *nanog* (MZn*anog*), two studies further proved that zebrafish *nanog* is primarily required for extraembryonic development [41] and it is crucial for embryonic architecture formation and cell survival [42].

In the present study, we independently generated two MZ*nanog* alleles with zebrafish, and demonstrated that maternal Nanog interacts with maternally deposited TCF, thereby safeguards the embryo against formation of β-catenin/TCF transcriptional activation complex which may induce hyper-dorsalization of the embryo. Our study thus uncovers a negative regulation system of maternal β-catenin activity, by revealing Nanog as a novel transcriptional switch between Nanog/TCF and β-catenin/TCF complexes.

## Results

### Tle3a and Tle3b do not contribute to the repression of maternal β-catenin activity

In zebrafish early embryonic development, maternal β-catenin proteins is generally believed to be translocated to the nuclei of dorsal blastomeres and subsequently activate the expression of a series of zygotic genes for dorsal commitment [43–47]. In a screening for maternal factors, we happened to observe that the nuclear β-catenin localized not only at the dorsal-most blastomeres with high amount but also in some ventral and lateral cells with low amount in early zebrafish embryos at 3.5 hour-post-fertilization (hpf) (Fig. 1a), just like that in *Xenopus*, lower levels of nuclear β-catenin also presents throughout the embryo at blastulae stage [48, 49]. Furthermore, we showed that overexpression of *β-catenin* at non-dorsal cells could induce ectopic expression of maternal β-catenin targets, *boz* and *chd* (Fig. S1), suggesting that the low level of ventrally located endogenous nuclear β-catenin activities should be repressed in early embryo.

**Figure 1.**
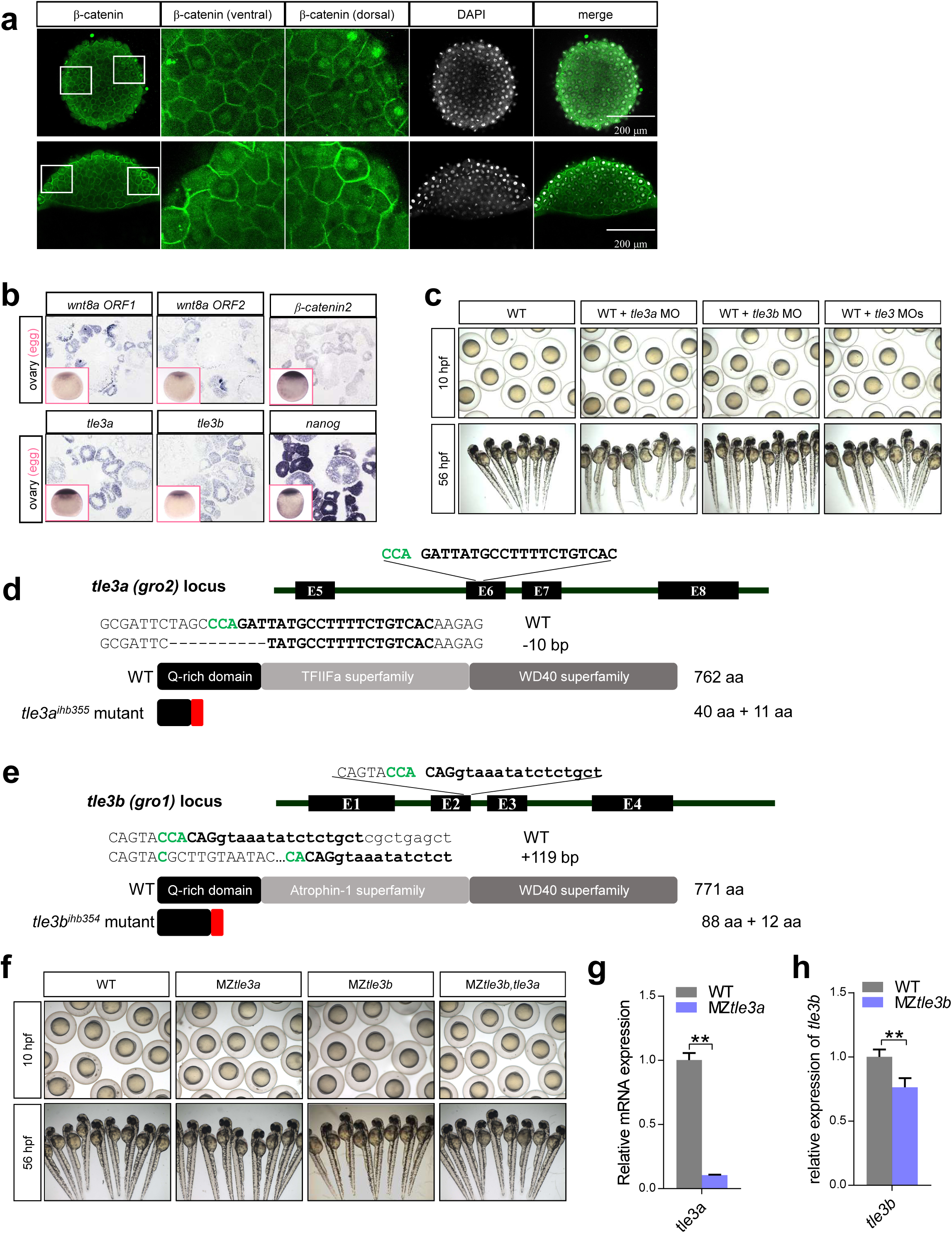
Tle3a and Tle3b do not contribute to the repression of maternal β-catenin activity. **(a)** Detection of nuclear localization of maternal β-catenin at 3.5 hpf by immunostainning. Upper, animal view. Lower, lateral view. Total-β-catenin was co-stained with DAPI. **(b)** *wnt8a1*, *wnt8a2*, *β-catenin*, *tle3a*, *tle3b* and *nanog* are maternally expressed during oogenesis and unfertilized eggs. Results showed cryosection and whole-mount *in situ* hybridization. **(c)** Injection of *tle3a* MO, *tle3b* MO, or *tle3* MOs together, does not lead to early developmental defect. MO was injected at one-cell stage and at least 50 embryos were injected and observed. **(d)** The CRISPR/Cas9 target of *tle3a* is located within exon 6, and a 10-bp deletion mutant (*tle3a^ihb355^*) was obtained. **(e)** The CRISPR/Cas9 target of *tle3b* is located at the splicing site of exon 2 and intron 2, and a 119-bp insertion mutant (*tle3b^ihb354^*) was obtained. Target sequence is in bold and PAM sequence is in Green. **(f)** The maternal and zygotic mutant of *tle3a* or *tle3b*, or double mutant of *tle3a* and *tle3b*, showed no early developmental defect. **(g)** The mRNA expression level of *tle3a* was reduced in MZ*tle3a* at 3hpf. **(h)** The mRNA expression level of *tle3b* was reduced in MZ*tle3b* at 3hpf. ** means p<0.01.

Therefore, we were curious about how the ventrally distributed nuclear β-catenin activities are controlled. One possibility is the presence of certain antagonistic factors of β-catenin inside nucleus, e.g., Groucho (Gro/TLE), which binds to repressive T-cell factor (TCF) and further interacts with histone deacetylases to maintain the chromatin in a transcriptionally inactive state [8–11, 50]. We carefully examined the expression of several Wnt/β-catenin signaling components - *wnt8* (*wnt8a ORF1* and *wnt8a ORF2*), *β-catenin 2* (*ctnnb2*), *tle3a* (*groucho2*) and *tle3b* (*groucho1*) during oogenesis and in the unfertilized eggs by *in situ* hybridization. Both *ctnnb2* and *wnt8a* were shown to be maternally expressed in the developing oocytes and the unfertilized eggs (Fig. 1b), further supporting that certain amount of β-catenin might enter the nuclei of all of the blastoderm cells and its transcriptional activity should be repressed in the non-dorsal most cells. Since both of *tle3a* and *tle3b* mRNA are maternally deposited (Fig. 1b), we performed loss-of-function studies by knockdown or knockout of *tle3a* and *tle3b*. To our surprise, either single or combined knockdown of *tle3a* or *tle3b* did not interrupt early embryogenesis (Fig. 1c). We further generated the maternal-zygotic mutants of *tle3a* and *tle3b* by CRISPR/Cas9 technology (Fig. 1d, e). Consistent to the knockdown experiments, either the single or double maternal-zygotic mutants of *tle3a* and *tle3b* did not show any defects of early embryonic development (Fig. 1f-h). All these data indicate that Tle3a and Tle3b do not contribute to the repression of maternal β-catenin activity and there might be other factors protecting the embryos from the activation of maternal Wnt/β-catenin activity at ventral region.

### Knockdown of *nanog* leads to dorsalization or posteriorization

During oogenesis, *nanog* is maternally expressed in the oocytes at different stages (Fig. 1b). After fertilization, *nanog* mRNA is ubiquitously distributed in the blastoderm cells till 30% epiboly, dramatically decreased at shield stage and becomes undetectable from 75% epiboly (Fig. 2a), implying a major role of zebrafish *nanog* during early development. To examine the function of *nanog* in early embryogenesis, we first utilized a previously published antisense morpholino (MO) targeting the translational start site of *nanog* mRNA to knock down *nanog*) [39]). Intriguingly, injection of low-dose (LD) *nanog* MO (0.5 ng per embryo) mainly led to forebrain defects (Fig. 2b), mimicking the zygotic overexpression of *wnt8a* [51]. We then increased the *nanog* MO dosage to 1.2 ng per embryo (moderate-dose, MD), and found that most of the embryos were strongly dorsalized (Fig. 2b), resembling the observation in a recent study [52].

**Figure 2.**
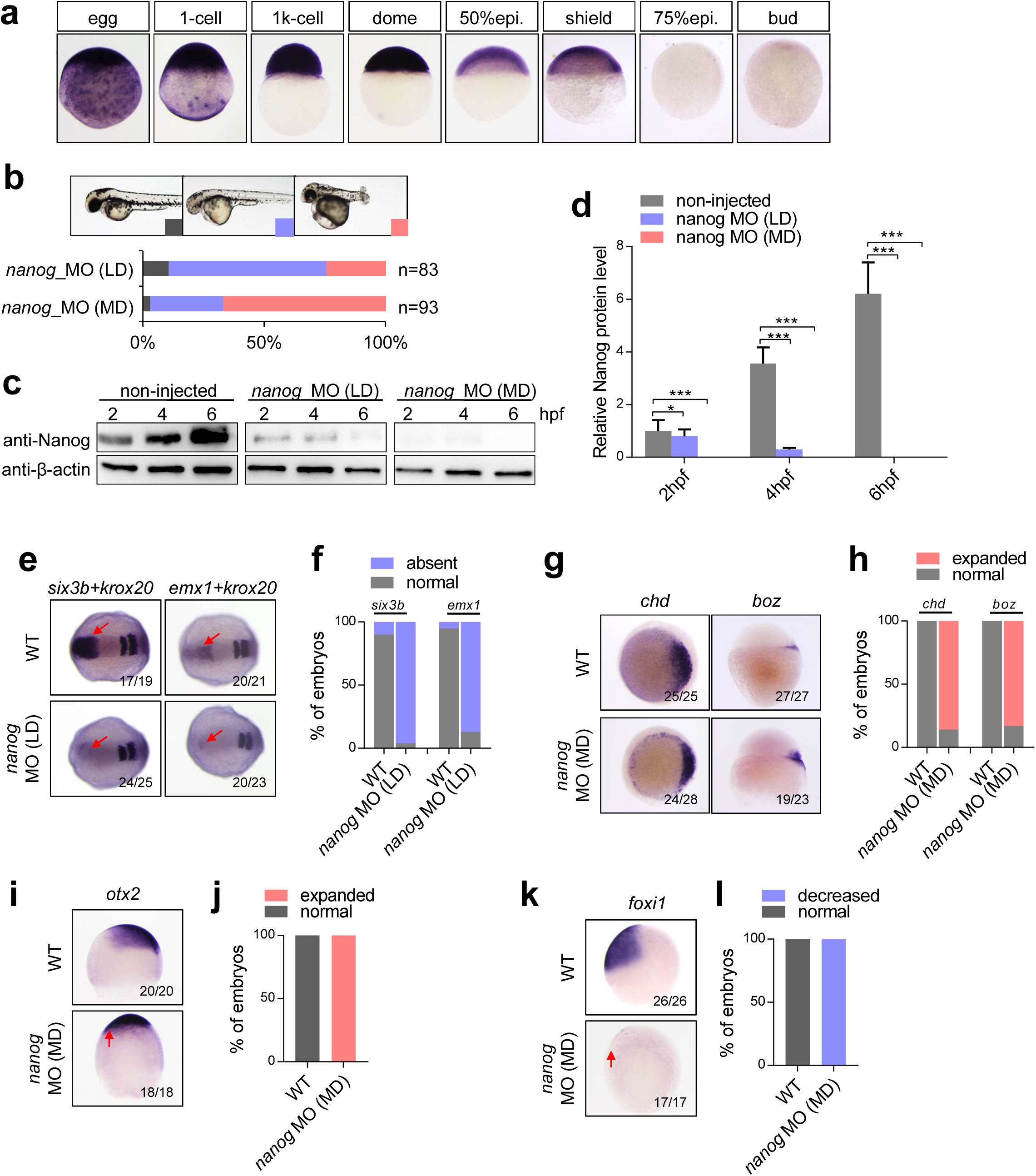
Knockdown of *nanog* leads to dorsalization and posteriorization. **(a)** *nanog* mRNA is maternally transcribed and vanishes at 75% epiboly stage. **(b)** Two different phenotypes are observed at two doses of *nanog* MO (0.5 ng/embryo; low dose, LD, and 1.2 ng/embryo, moderate dose, MD) injected embryos, telencephalon defect and dorsalization. Phenotype was observed at 36 hpf. n represents embryo numbers. **(c)** Western blot detection of Nanog in two doses of *nanog* MO injected embryos. Nanog translation was blocked in all detected stages in moderate dose of *nanog* MO injected embryos, and few of Nanog protein can be detected at early stage in low dose of *nanog* MO injected embryos. **(d)** Relative Nanog signal intensity in western blots. **(e)** and **(f)** The expressions of forebrain marker *six3b* and telencephalon marker *emx1* are absent in low dose of MO injected embryos. *krox20* was used as a stage-control marker. Red arrows indicate the expression region of *six3b* or *emx1*. (**g**) and **(h)** The dorsal marker gene, *chd*, and maternal β-catenin target, *boz*, were both up-regulated in moderate dose of *nanog* MO injected embryos. **(i)** and (**j**) The expression of dorsal neuroectoderm marker *otx2*, is expanded in low dose of *nanog* MO injected embryos. Red arrow indicates the ventral expansion of *otx2*. **(k)** and **(l)** The ventral epidermal ectoderm marker *foxi1* is eliminated in low dose of *nanog* MO injected embryos. Red arrow indicates the ventral absence of *foxi1*. *foxi1* and *otx2* were detected at 90% epiboly stage, *six3b* and *emx1* were detected at 2-somite stage. *chd* was detected at 4.5 hpf, *boz* was detected at 4 hpf. *** means p<0.001, * means p<0.05.

To clarify the different phenotypes after injection of different dosages of *nanog* MO, we performed Western blot to examine the Nanog protein levels at different stages in wildtype (WT) and the embryos (morphants) injected with low-dose or moderate-dose of MO. As shown in Figure 2c and 2d, injection of low-dose *nanog* MO only led to partial reduction of Nanog protein expression level at 2 hour-post-fertilization (hpf) and complete elimination of Nanog protein at 6 hpf, indicating that the endogenous Nanog proteins was eliminated after the time of zygotic genome activation (ZGA). Whereas a moderate-dose of *nanog* MO led to complete absence of Nanog protein from 2 hpf (Fig. 2c, d), indicating a block of the maternal activity of Nanog proteins. Whole-mount *in situ* hybridization (WISH) analysis further confirmed the different phenotypes. After injection of low-dose *nanog* MO, the expression of forebrain marker *six3b* and telencephalon marker *emx1* were nearly absent (Fig. 2e, f), mimicking the zygotic activation of Wnt/β-catenin signaling [18, 53]. However, in the moderate-dose *nanog* morphants, the expression levels and expression territories of maternal β-catenin signaling targets *boz* and *chd* were strongly increased (Fig. 2g, h), mimicking the dorsalization phenotype resulting from the ectopic activation of maternal β-catenin signaling (Supplemental Fig. S1, [18]). At mid-gastrulation, the moderate-dose morphants showed strong expansion of *otx2* (dorsal ectoderm marker) (Fig. 2i, j) and ventral shrinkage of *foxil* (ventral ectoderm marker) (Fig. 2k, l), further demonstrating the dorsalization phenotype in the moderate-dose *nanog* morphants. Taken together, the above results suggest that elimination of Nanog activities before ZGA or after ZGA gave different phenotypes, mimicking the maternal or zygotic activation of Wnt/β-catenin signaling, respectively.

### Knockdown of *nanog* activates Wnt/β-catenin signaling

To further study the relationship between Nanog and Wnt/β-catenin signaling pathway, we performed a series of genetic interaction experiments. First, we compared the low-dosage *nanog* morphants with the zygotic Wnt//β-catenin signaling activated embryos. The low-dosage *nanog* morphants have no eyes or telencephalon, similar to *wnt8a* (1 pg per embryo) overexpressed embryos or the *tcf7l1a* (previous name: *tcf3a* or *headless*) depleted embryos (*tcf7l1a* MO, 1.6 ng per embryo) (Fig. 3a) [18, 53]. We then titrated the dosages of *nanog* MO (160pg per embryo), *wnt8a* mRNA (0.1pg per embryo) and *tcf7l1a* MO (800pg per embryo) to obtain normal brain patterning in the injected embryos. However, when *nanog* MO was co-injected with *wnt8a* mRNA or *tcfl1a* MO, a majority of embryos developed with the headless phenotype (Fig. 3b), suggesting the genetic interaction between *nanog* and zygotic Wnt signaling. To further investigate the crosstalk between Nanog and Wnt/β-catenin signaling, we performed genetic interaction experiments. When compared with WT, the expression of zygotic Wnt target gene *sp5l* was expanded to the animal pole and ventral ectoderm, and the expression of Wnt antagonist *dkk1b* and *frzb* was decreased in the *nanog* morphants (Fig. 3c, d). Injection of *wnt8a* MO largely restored the expression of those genes in *nanog* morphants (Fig. 3c, d). More strikingly, the expression of two telencephalon markers – *six3b* and *emx1*, which were absent in the low-dose *nanog* MO injected embryos, were recovered after *wnt8a* knockdown (Fig. 3c, d). All these data indicate that Nanog negatively regulates Wnt/β-catenin signaling.

**Figure 3.**
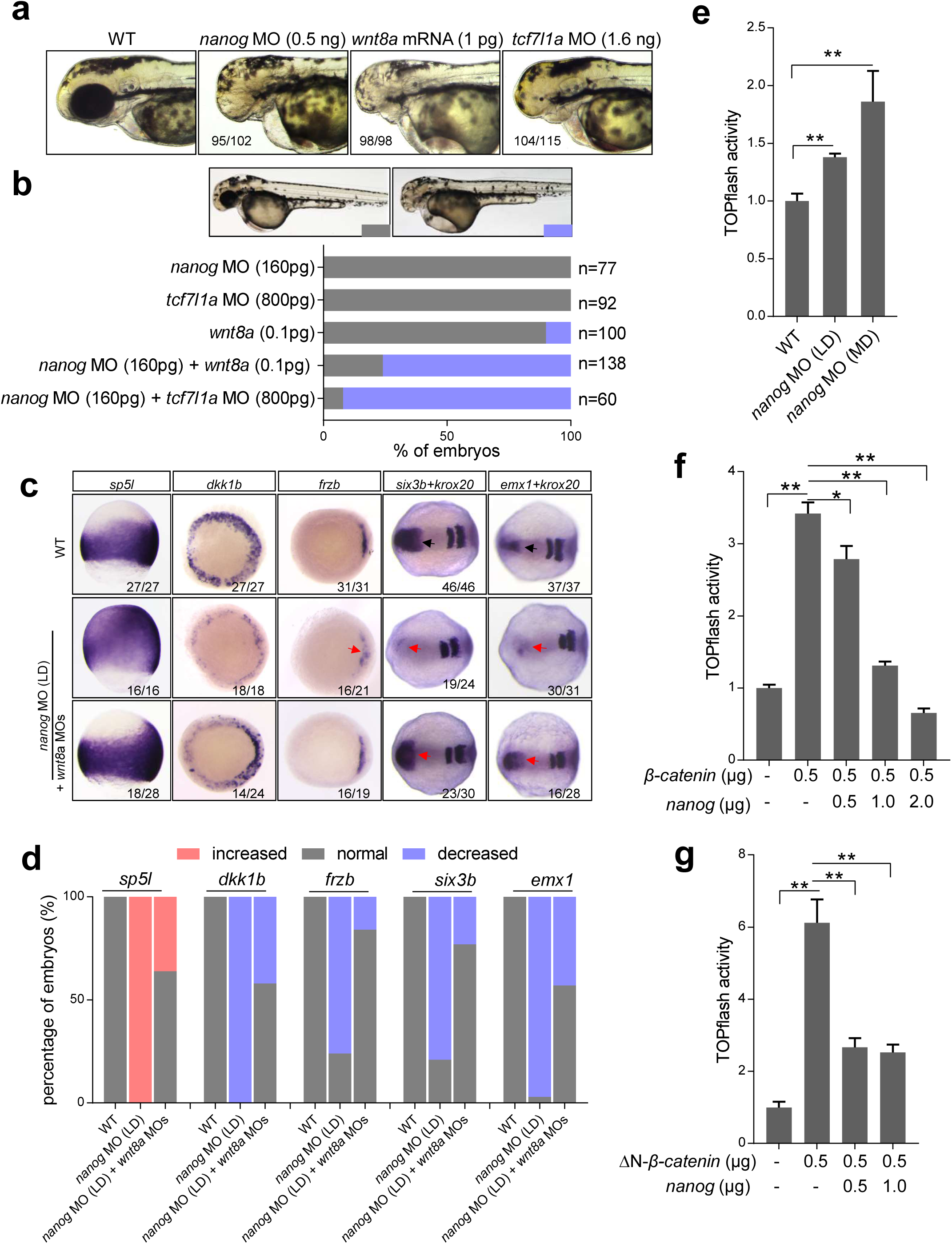
*nanog* negatively regulates Wnt/β-catenin signaling. (**a**) The embryos injected with low dose of *nanog* MO (0.5 ng) exhibit the similar phenotypes-telencephalon defect - with *wnt8a* (1 pg) overexpression embryos and *tcf7l1a* (1.6 ng) knockdown embryos. Phenotype was observed at 36 hpf. (**b**) Embryos injected with *nanog* MO (160 pg), *wnt8a* mRNA (0.1 pg) or *tcf7l1a* MO (800 pg) at extremely low dose show no obvious defect, respectively. However, co-injection of *nanog* MO with *wnt8a* mRNA or *tcf7l1a* MO at the same dose resulted in telencephalon truncated. n represent the number of embryos we observed. (**c**) Wnt/β-catenin activity is over-activated in *nanog* morphants and can be rescued by knockdown of *wnt8a*. The expression of zygotic Wnt target genes, *sp5l* and *frzb* were up-regulated in *nanog* morphant whereas Wnt antagonist *dkk1b* was reduced, and this expression defect can be restored by knockdown of *wnt8a*. so did the telencephalon defect in *nanog* morphant. *gfp* mRNA was used as injection control. *sp5l* was detected at 75% epiboly, *frzb* and *dkk1b* were detected at shield stage, *six3b* and *emx1* were detected at 2-somite stage, *krox20* was used as stage control. (**d**) The percentage of embryos counted in WISH **(c)**. (**e**) Analysis of TOPflash activity showed Wnt/β-catenin activity is up-regulated in both of low dose and moderate dose of *nanog* MO injected embryos. Embryos were collected at 8 hpf. (**e**) Analysis of TOPflash activity showed co-transfection of Nanog with β-catenin in HEK293T cells inhibited the up-regulated Wnt/β-catenin activity induced by β-catenin in a dose-dependent manner. (**f**) Overexpression of *nanog* in HEK293T cells can also inhibit the up-regulated Wnt/β-catenin activity induced by ΔN-β-catenin (constitutively activated β-catenin). ** means p<0.01, * means p<0.05.

We then investigated the repressive role of Nanog on Wnt/β-catenin signaling activity by *in vivo* and *in vitro* TOPflash assay. Knockdown of *nanog* with low and moderate dosages of *nanog* MO resulted in dose-dependent up-regulation of Wnt signaling activity in embryos (Fig. 3e). In 293T cells, the TOPflash activity was significantly increased after transfection of WT β-catenin. However, when the cells were co-transfected with different amounts of Nanog, the TopFlash activity showed significantly reduction in a dose-dependent manner (Fig. 3f). We then performed a similar TopFlash assay in the cells transfected with ΔN-β-catenin, which can sustainably enter the nucleus to active the transcriptional activity [29]. To our surprise, co-transfection of Nanog could still significantly repress the Wnt activity resulting from overexpression of “activated form” of β-catenin (ΔN-β-catenin) in a dose dependent manner (Fig. 3g). These results indicate that the Nanog could effectively repress the transcriptional activity of nucleus-located β-catenin.

### Maternal β-catenin activity is hyper-activated in MZ*nanog* embryo

In order to fully characterize the role of *nanog* in early embryonic development, we generate *nanog* mutants using TALEN mediated mutagenesis as described in our previous study [54]. We identified two mutant lines - one with 2-bp deletion (−2) and the other one with 1-bp insertion (+1) (Fig. S2a). The 2-bp deletion resulted in the frame-shift of the open reading frame. The 1-bp insertion resulted in premature termination at the target site and encoding a truncated Nanog of 18 amino acids (Fig. S2a). The two alleles were named as *nanog^ihb97/ihb97^* (2-bp deletion) and *nanog^ihb98/ihb98^* (1-bp insertion). After in-cross with the *nanog* heterozygous, we found that the zygotic mutant of *nanog* (Z*nanog^ihb97^*) did not show any embryonic defect and grew up to adulthood (Fig. S2b). We then continued to in-cross the zygotic mutant and obtained the maternal zygotic mutant of *nanog* (MZ*nanog*). Compared with WT and Z*nanog* embryos, the MZ*nanog^ihb97^* failed epiboly movement and all cell stacked at the animal pole. MZ*nanog^ihb98^* exhibited less severe phenotype. Most of the embryos from both alleles of MZ*nanog* could not survive to 24 hpf (Fig S2b). We then obtained maternal mutant of *nanog* (M*nanog^ihb97^*) by mating MZ*nanog^ihb97^* female with WT male. M*nanog^ihb97^* embryos are phenotypically identical to MZ*nanog^ihb97^*, implying that the early development was mainly regulated by maternal deposit *nanog* mRNA (Fig. S2b).

To verify that the Nanog protein was completely absent in MZ*nanog*, we detected the protein level of Nanog by western blot. In WT embryos, Nanog protein could be detected from 64-cell stage and decreased rapidly during gastrulation. After the 75% epiboly stage, Nanog protein was no longer detected, whereas the Nanog protein could not be detected in both types of MZ*nanog* embryos (Fig. 4a). Immunostaining analysis of Nanog further confirmed that the Nanog is mainly localized in the cell nucleus and there is no Nanog expression in the MZ*nanog* (Fig. 4b). These data suggested that the Nanog was completed disrupted in MZ*nanog* mutant.

**Figure 4.**
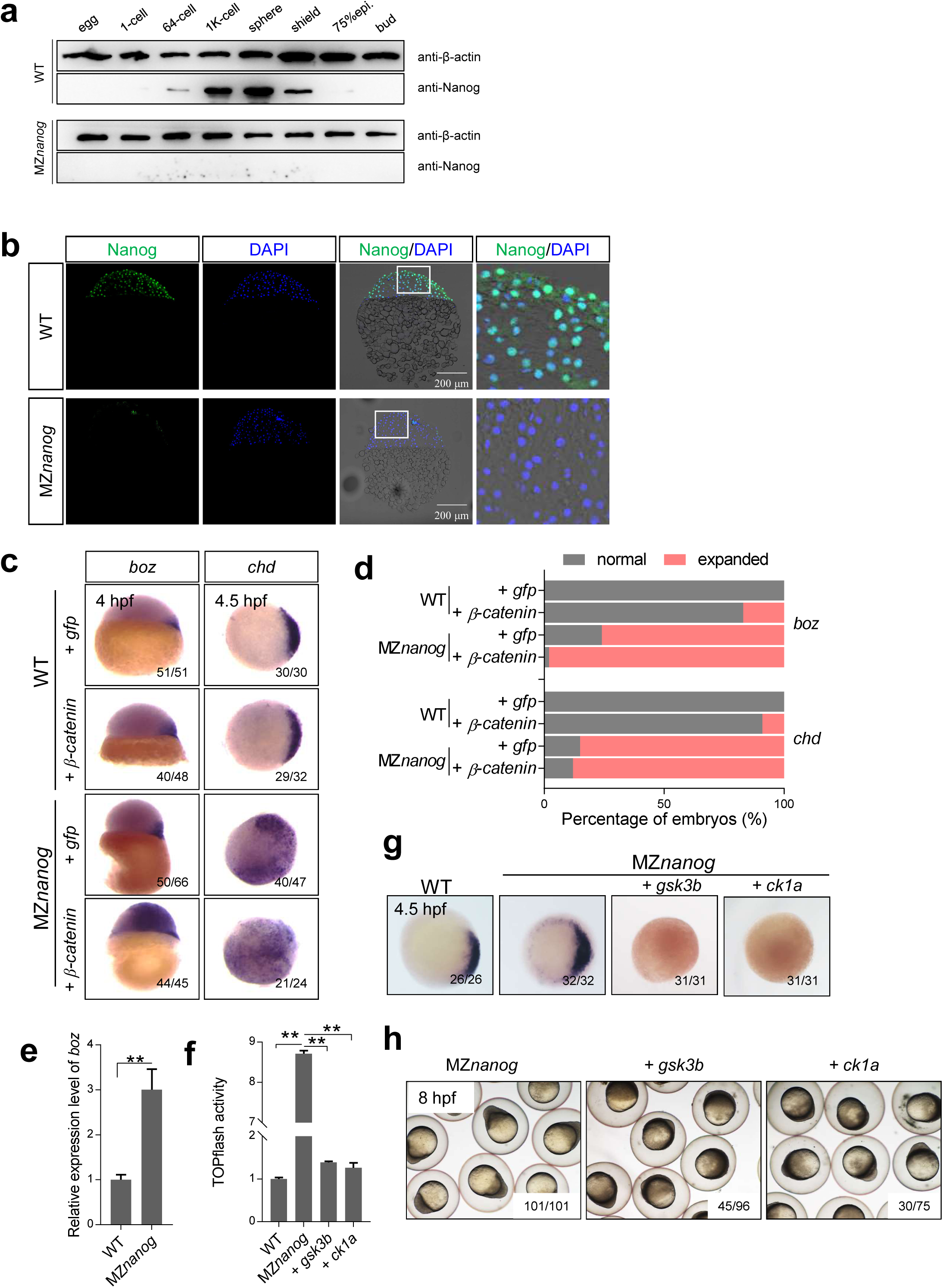
Wnt/β-catenin activity is hyper-activated in *nanog* mutant. (**a**) Translation of Nanog protein totally disappeared in MZ*nanog*. Nanog protein can be detected as early as 64-cell stage and vanished at 75% epiboly stage in WT embryos, while no Nanog protein was detected in MZ*nanog*. (**b**) Immunolocalization of Nanog on cryosections of WT and MZ*nanog* at sphere stage. Nanog is localized in cell nucleus of WT and disappeared in MZ*nanog*. (**c**) Injection of low dose of β-catenin mRNA (200 pg) induced slightly up-regulation of *boz* and *chd* in WT embryos, but highly increasing in MZ*nanog* embryos, indicating that Wnt activity is over-activated because of *nanog* LOF. Injection or co-injection was performed at one-cell stage. *boz* was detected at 4 hpf, *chd* was detected at 4.5 hpf. (**d**) The percentage of embryos counted in WISH of **(c)**. (**e**) RT-qPCR showed the expression of *boz* is significantly up-regulated in MZ*nanog*. Embryos were collected at 4 hpf. (**f**) TOPflash activity assay showed that Wnt/β-catenin signaling activity was significantly up-regulated in MZ*nanog* embryos, and could be restored by overexpression of Wnt antagonists, *gsk3b* or *ck1a*. (**g**) Ectopic expression of *chd* in MZ*nanog* was also rescued by *gsk3b* and *ck1a* overexpression. *chd* was detected at 4.5 hpf. (**h**) Overexpression of *gsk3b* or *ck1a* was confirmed to rescue the early development defect of MZ*nanog*. Embryos were injected at one-cell stage and phenotype was observed at 8 hpf. ** means p<0.01, 3 biological replicates were performed, student’s *t-test*.

We also compared the transcript level of *nanog* in MZ*nanog^ihb97^* and MZ*nanog^ihb98^*. The maternal *nanog* mRNAs were absent in the MZ*nanog^ihb97^*. Zygotic transcription of *nanog* appeared to happen at 1000-cell but quickly vanished at 30% epiboly (Fig. S2c). The mutated *nanog* mRNA expressed from 2-cell stage until 50% epiboly but disappeared at 90% epiboly in MZ*nanog^ihb98^* (Fig. S2c). Due to the slight presence of mutated *nanog* transcripts in MZ*nanog^ihb98^*, we used MZ*nanog^ihb97^* (for abbreviation: MZ*nanog*) for the subsequent experiments.

Just like the *nanog* morphants, MZ*nanog* showed strong activation of maternal β-catenin activity, as indicated by expanded expression region of *boz* and *chd* (Fig. 4c, d). RT-qPCR also confirmed that the *boz* expression was remarkably up-regulated in MZ*nanog* in comparison with WT (Fig. 4e). When we injected low-dose of β-catenin mRNA (100 pg per embryo) into WT or MZ*nanog* embryos, the MZ*nanog* showed a dramatic expansion of *boz* and *chd*, although low dose of β-catenin could slightly increase the expression of *boz* and *chd* in WT embryos (Fig. 4c, d). TopFlash assay further supported that the Wnt activity was elevated in MZ*nanog* embryos, and this effect could be suppressed by overexpression of *gsk3b* or *ck1a* (Fig. 4f). WISH analysis of *chd* at sphere stage further confirmed the rescue effect by *gsk3b* or *ck1a* (Fig. 4g). More interestingly, by overexpression of *gsk3b* or *ck1a*, the hyper-dorsalization defects of MZ*nanog* at gastrula stage could be partially rescued (Fig. 4h). These results illustrate that loss of *nanog* function leads to over-activation of maternal Wnt/β-catenin activity.

### Depletion of Nanog does not affect the nuclear translocation of β-catenin

To determine whether the over-activation of Wnt/β-catenin activity in MZ*nanog* was caused by the upregulation of maternal *wnt8a* or *β-catenin2*, we examined the expression of those transcripts. By WISH analysis, we showed that the expression of *wnt8a* (*wnt8a ORF1* and *wnt8a ORF2*) and *β-catenin2* appeared comparable between WT and MZ*nanog* in unfertilized egg and in 16-cell stage embryos (Fig. 5a). By RT-qPCR analysis, we found that *wnt8aORF1* even showed significantly decreased expression level when compared with WT in the ovary, and embryos at 1-cell, 2 hpf and 4 hpf (Fig. 5b and c). These data indicate that the hyperactivation of maternal Wnt activity in MZ*nanog* is not due to the up-regulation of genes encoding Wnt ligands or β-catenin.

**Figure 5.**
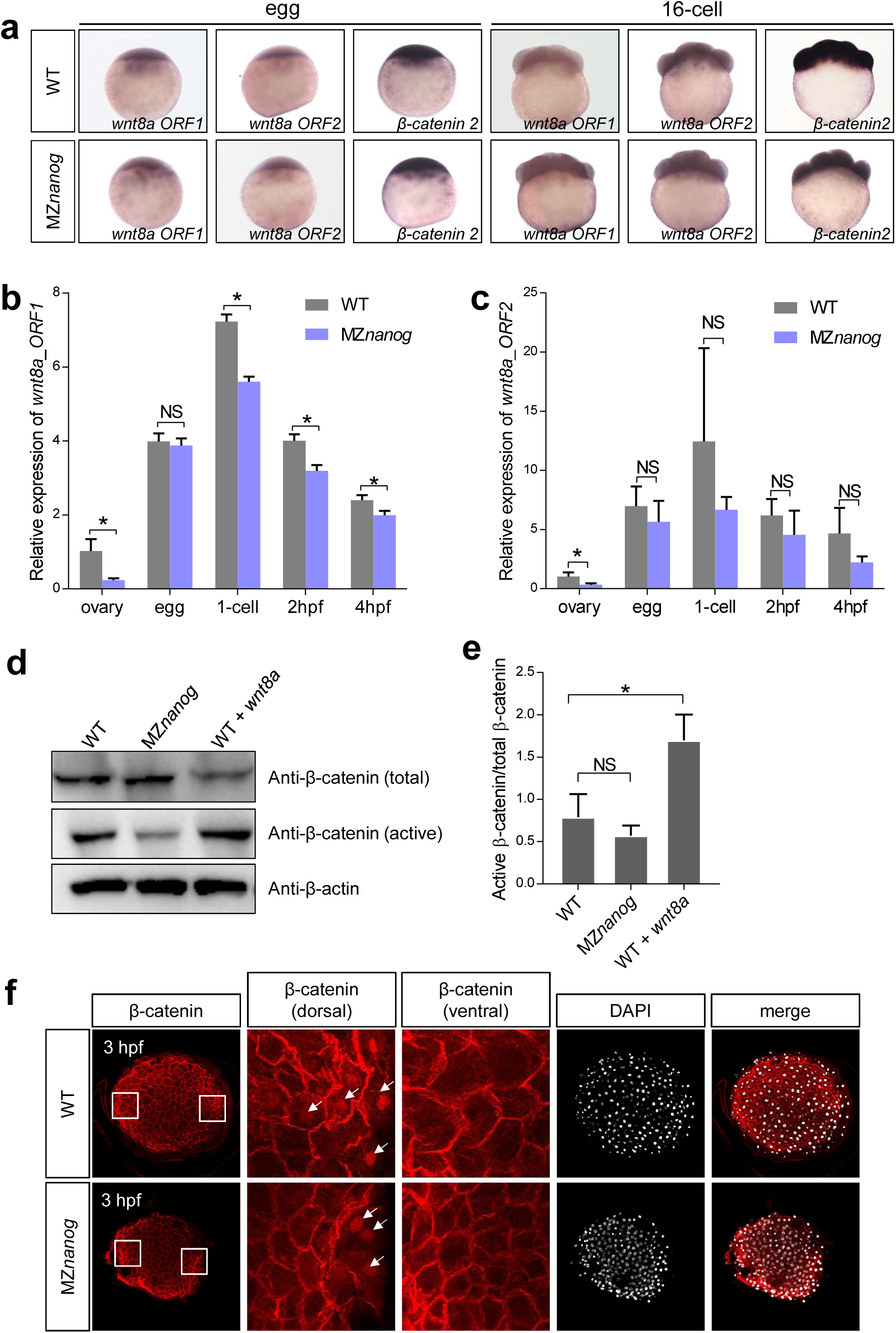
Nanog does not regulate the transcription of Wnt/β-catenin components and nuclei localization of β-catenin. (**a**) In situ hybridization and (**b, c**) RT-PCR showed the maternal transcription of *wnt8a1*, *wnt8a2*, and *β-catenin2* were not affected in MZ*nanog* when compared with WT. (**d**) Western blot and (**e**) statistical results show that the level of nuclear β-catenin (anti-active β-catenin) in MZ*nanog* was not increased. Anti-total β-catenin was used as β-catenin expression control and anti-β-actin was used as internal control. 2 pg of *wnt8a* mRNA was injected at one-cell stage in WT and used as positive control. Embryos were collected at 4 hpf. Experiments were carried out for triplicates. (**f**) Immunolocalization of β-catenin on whole-mount embryos at 1000-cell stage (3hpf) shows that the nuclear β-catenin in both of WT and MZ*nanog* was localized in dorsal margin cells and devoid of ventral cells, indicating that Nanog does not regulate the nuclei localization of β-catenin. Signal was observed at animal view. NS, no significant difference. * means p<0.05.

It is well-known that the activation of canonical Wnt pathway leads to translocation of β-catenin into the cell nucleus [13, 55]. We then aimed to test whether the over-activation of Wnt activity in MZ*nanog* is caused by increased accumulation of β-catenin in the nucleus. Firstly, we examined the level of nuclear β-catenin in MZ*nanog* at 4 hpf by western blot by using *wnt8a*-overexpressed embryo as positive control. Unlike that the ratio of active β-catenin/total β-catenin was remarkably increased in *wnt8a*-overexpressed embryos, there was no significant difference between MZ*nanog* and WT embryos (Fig. 5d, e). We further checked the nuclear β-catenin accumulation in MZ*nanog* by immunostaining at 3 hpf, and found that the nuclear localized β-catenin in MZ*nanog* appeared comparable to that in WT (Fig. 5f). Taken together, we demonstrate that the hyper-activation of maternal β-catenin activity is not due to the increased nuclear translocation of β-catenin.

### N-terminal of Nanog is required for suppression of Wnt/β-catenin signaling

As Nanog is a type of homeobox protein, we investigated whether Nanog functions as a transcriptional activator or repressor. We fused the homeodomain of Nanog with the transcriptional activator domain Vp16 or the transcriptional repressor domain of Engrailed 2 [56, 57], to generate overexpression constructs of constitutive-activator type (Vp16-Nanog) or constitutive-repressor type (En-Nanog) of Nanog (Fig. S3a). After injection of *vp16-nanog* mRNA into WT embryos, transcriptional targets of Nanog at zygotic genome activation (ZGA), *mxtx2*, *blf* and *mir-430*, showed increased expression, mimicking overexpression of WT *nanog* (Fig. S3b, c). In contrast, injection of ENH mRNA led to decreased expression of those genes, mimicking *nanog* morphants (Fig. S3b, c). On the other hand, the expression of *sod1*, a maternal mRNA targeted by miR-430 [58], was accumulated in *nanog* morphants and the En-Nanog overexpressed embryos, indicating the defects of maternal mRNA clearance (Fig. S3b, c). Further GFP-3xIPT-miR-430 reporter assay and RT-qPCR analysis of miR-430a and miR-430b confirmed the defects of maternal to zygotic transition (MZT, mainly containing the events of maternal mRNA clearance by miR-430 and ZGA) in MZ*nanog* and the rescue effects by overexpression of full-length Nanog (Nanog_FL), Vp16-Nanog and miR-430 (Fig. S3d-f). All these demonstrate that Nanog serves as a transcriptional activator during maternal to zygotic transition (MZT), by activating the zygotic genes and miR-430 which can further clean the maternal mRNAs.

To understand the transcriptional activation effects of Nanog, we injected full-length *nanog*_FL mRNA, *vp16-nanog* mRNA and C-terminal truncated *nanog* (*nanog*_ΔCT) mRNA into MZ*nanog* embryos, to compare their rescue effects. WISH analysis of *mxtx2*, *blf*, *mir-430* and *sod1* proved the defects of yolk syncytial layer (YSL) and defective MZT in MZ*nanog*, and overexpression of Nanog_FL, Nanog_ΔCT, or Vp16-Nanog could successfully rescue the defects of YSL development and MZT in MZ*nanog* (Fig. 6a, b), which further supported the notion that Nanog functions as transcriptional activator during MZT. Nevertheless, when we checked the overall development of the embryos, overexpression of either Nanog_FL or Nanog_ΔCT could fully rescue the mutant phenotype and the embryos could even survive to adulthood (Fig. 6c, d, e), indicating that the C-terminal of Nanog is not required for its normal function. On the other hand, although overexpression of *vp16-nanog* nearly rescued all the developmental defects of MZ*nanog*, the rescued embryo still showed a head-truncation phenotype (Fig. 6c, d, e), indicating that the elevated Wnt/β-catenin activity existing in the rescued embryos. Moreover, unlike overexpression of Nanog_FL or Nanog_ΔCT which could efficiently suppress β-catenin-induced TOPFlash activity, overexpression of Vp16-Nanog could not inhibit the TOPFlash activity (Fig. 6f). All these strongly suggest that N-terminal of Nanog is required for its suppression activity on Wnt/β-catenin signaling.

**Figure 6.**
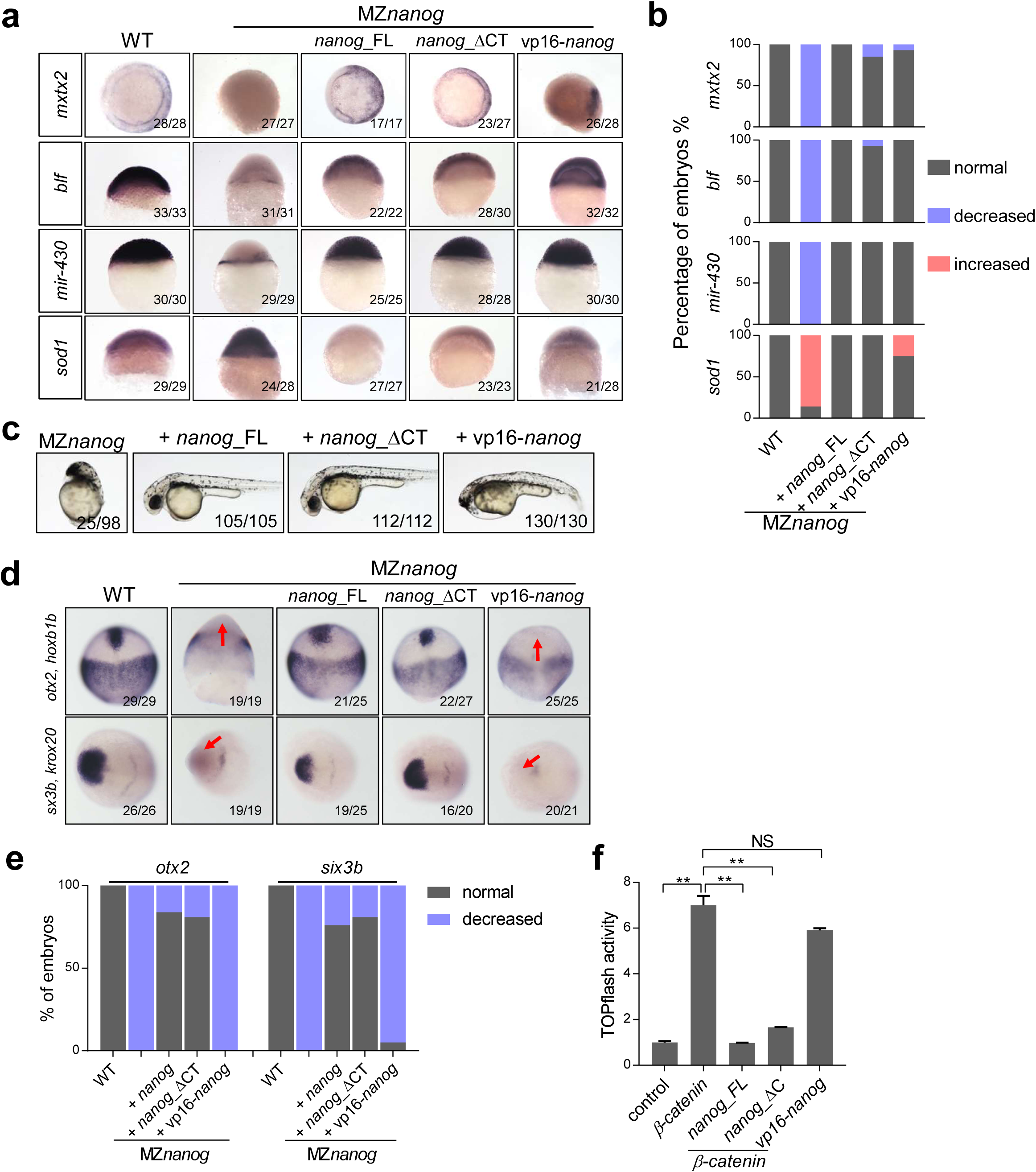
N-terminal of Nanog is required for its Wnt/β-catenin repressive activity. (**a**) The expression of mesendoderm marker, *mxtx2*, strictly zygotic gene, *blf*, and microRNA-430 precursor (*mir-430*) were reduced, even absent in MZ*nanog* embryos, while miR-430 target, *sod1* was significantly increased in MZ*nanog*. Overexpression of *nanog_*FL, *nanog*_ΔCT, or *vp16*-*nanog* can restore the expression of *mxtx2*, *blf*, *sod1* and *mir-430* in MZ*nanog*. *mxtx2*, *blf* and *sod1* were detected at shield stage, *mir-430* was detected at 4 hpf. (**b**) The percentage of embryos counted in WISH of **(a)**. (**c**) Overexpression of full length of *nanog* (*nanog_*FL), C-terminal truncated Nanog (*nanog*_ΔCT), and vp16-*nanog* homeodomain (*vp16*-*nanog*) can rescue the developmental defects of MZ*nanog*. Both of *nanog_*FL and *nanog*_ΔCT rescued embryos showed WT-like phenotype, and could grow up and reproduce offspring, while *vp16*-*nanog* rescued embryos still shows a phenotype of head truncation. Phenotype was observed at 36 hpf. (**d**) The expression of neuroectoderm marker, *otx2*, and forebrain marker *six3b* were absent in MZ*nanog* embryos, and can be restored by overexpression of *nanog_*FL or *nanog*_ΔCT, but not *vp16*-*nanog*. (**e**) The percentage of embryos counted in WISH of **(d)**. (**f**) TOPflash activity showed co-transfection of VP16-nanog with β-catenin in HEK293T cells could not inhibit the up-regulated Wnt/β-catenin activity induced by β-catenin.

### Nanog interferes with the binding of β-catenin to TCF7 *in vitro*

Since Nanog dose not regulate the nuclear β-catenin level and its suppression of β-catenin transcriptional activity relies on its N-terminal, we inquired whether Nanog physically interacts with β-catenin or its nuclear partners, such as TCF/Lef. To start with, we firstly identified that Tcf7 and Tcf4 are both activator-type TCF, since localized injection of *tcf7* or *tcf4* mRNA alone, or co-injection with *β-catenin* mRNA, *tcf7* and *tcf4* could efficiently induce the ectopic expression of maternal β-catenin targets, *boz* and *chd* (Fig. S4), supporting the previous report that Tcf7 acts as β-catenin-dependent trans-activators with Lef1 [59–61].

We then performed a co-immunoprecipitation (co-IP) assay, and found that Nanog physically interacts with Tcf7 (Fig. 7a). To determine that which part of Nanog is capable of binding with Tcf7, we generated serval deletion types of Nanog, including full-length *nanog* (myc-Nanog), N-terminal truncated Nanog (myc-Nanog-ΔN), homeodomain deleted Nanog (myc-Nanog-ΔH), C-terminal truncated Nanog (myc-Nanog-ΔC) and N-terminal-only Nanog (myc-Nanog-NT). In the co-immunoprecipitation screening, we found that all types of Nanog proteins except Nanog-ΔN bind to Tcf7 *in vitro* (Fig. 7a), suggesting the N-terminal of Nanog physically interacts with Tcf7.

**Figure 7.**
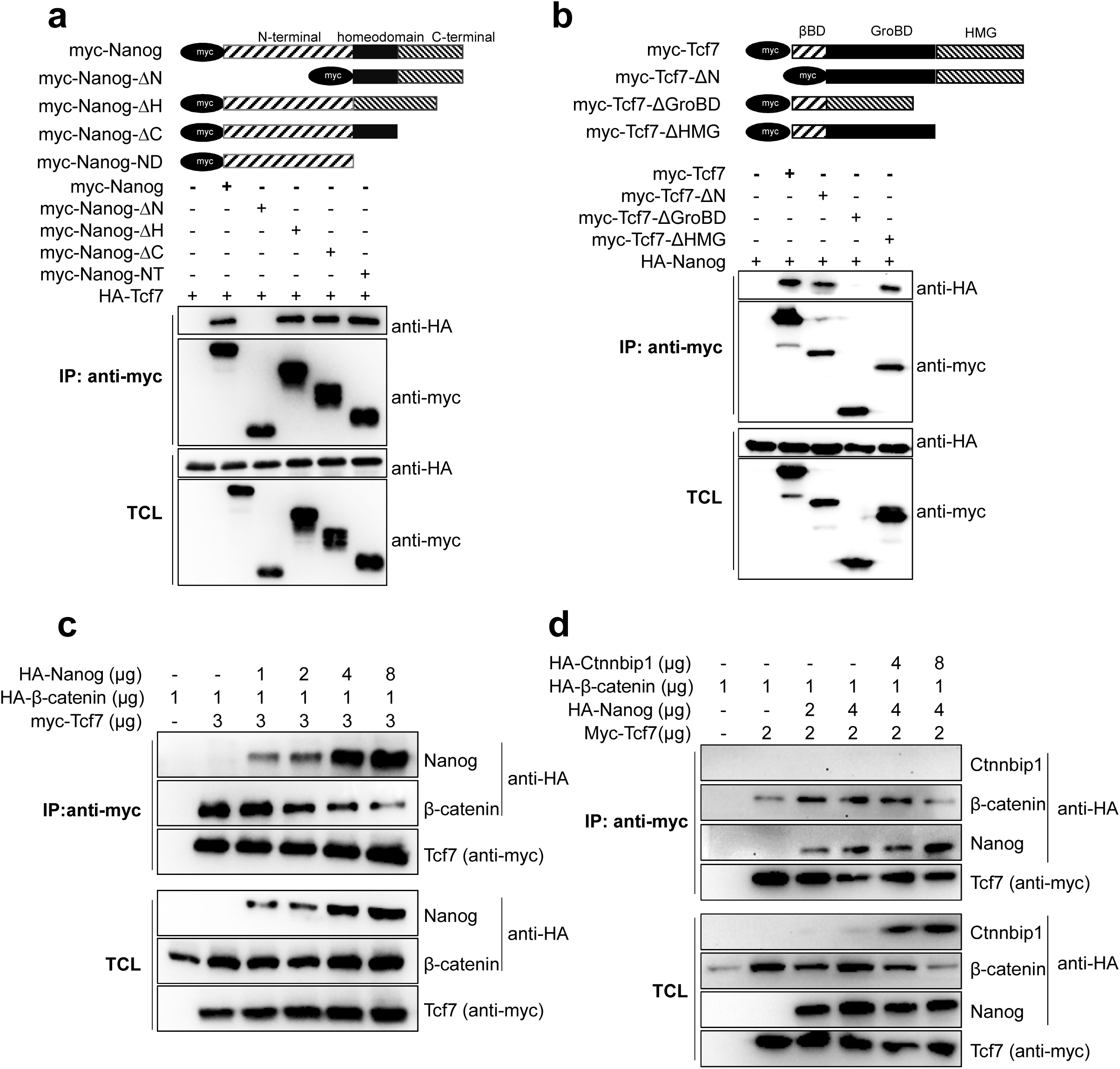
Nanog interferes with the binding of β-catenin to TCF7 *in vitro*. (**a**) Nanog interacts with TCF7 through its N-terminal. Different Myc-tagged Nanog were constructed and co-transfection with HA-Tcf7 in HEK293T cells. Deletion of Nanog N’ terminal could not coprecipitate with Tcf7, indicating that Nanog physically interacts with Tcf7 through N’ terminal. (**b**) TCF7 combines with Nanog through its Gro-binding domain. Different Myc-tagged Tcf7 were constructed and co-transfection with HA-Nanog in HEK293T cells. Deletion of Tcf7 Groucho-binding domain (GroBD) could not coprecipitate with Nanog, indicating that Tcf7 binds with Nanog through GroBD. (**c**) Nanog and β-catenin competitively binds with Tcf7. Co-transfection of increasing amount of Nanog decreases the interaction of β-catenin and TCF7 in a dose-dependent manner. More nanog was transfected into Tcf7 and β-catenin co-transfected cells, less β-catenin could be coprecipitated. The molecular weight of HA-Nanog is around 55 KDa and HA-β-catenin is around 100 KDa, so we can easily distinguish the anti-HA bands through protein size. (**d**) Co-transfected of Ctnnbip1 facilities the binding of Nanog with TCF7 and degradation of β-catenin. Two different amounts of HA-ctnnbip1 was co-transfected with Tcf7, Nanog and β-catenin in HEK293T cells, as the Ctnnbip1 expressed, more Nanog was coprecipitated and less β-catenin was pulled down. The molecular weight of HA-ctnnbip1 is around 10 KDa. Note that HA-β-catenin level is reduced when Ctnnbip1 is expressed.

Next, we aimed to understand that which part of Tcf7 is responsible for its interaction with Nanog. We generated different forms of Tcf7 including full-length Tcf7, β-catenin binding domain-deleted form (myc-Tcf7-ΔN), Groucho/TLE binding domain-deleted form (myc-Tcf7-ΔGroBD) and HMG deleted form (myc-Tcf7-ΔHMG). First, we confirmed the Groucho/TLE binding domain, β-catenin binding domain and Lef1 binding domain of Tcf7. As shown in Supplemental Figure S5, the full-length Tcf7 could efficiently bind to Gro2 (Fig. S5a), β-catenin (Fig. S5b) and Lef1 (Fig. S5c), whereas, the myc-Tcf7-ΔGroBD, myc-Tcf7-ΔN and myc-Tcf7-ΔHMG could not bind to Gro2, β-catenin and Lef1, respectively (Fig. S5a-c), which confirmed the respective binding domains of Gro2, β-catenin and Lef1 on Tcf7. We then conducted a similar co-IP assay to learn the Nanog binding domain of Tcf7. As shown in Figure 7b, all forms of Tcf7 except the Tcf7-ΔGroBD could efficiently interact with Nanog, illustrating that Nanog and Groucho/TLE both have high binding affinity to the Groucho/TLE binding domain of Tcf7. Taken together, the N-terminal region of Nanog physically interacted with the Groucho/TLE binding domain (GroBD) of Tcf7.

Since Nanog and β-catenin could both bind to Tcf7, there is a possibility that Nanog may interfere the interaction between β-catenin and TCF. We then performed a competitive binding assay in which the input of β-catenin and TCF was consistent in each sample and the input of Nanog was gradually increased. Strikingly, we found that the increased amount of Nanog effectively decreased the binding affinity of Tc7 to β-catenin (Fig. 7c), indicating that Nanog negatively regulate β-catenin transcriptional activity by attenuation of Tcf7/β-catenin transcriptional activator complex.

To challenge this conclusion, we co-transfected Ctnnbip1 (previously named ICAT), which can physically bind to β-catenin to prevent the association of β-catenin and TCF [28, 62, 63]. Interestingly, after co-transfected with Ctnnbip1, more Nanog was co-immunoprecipitated with Tcf7 (Fig. 7d), further supporting the conclusion that Nanog and β-catenin competitively interact with TCF7.

### Nanog interferes the β-catenin-TCF7 transcriptional complex *in vivo*

The above data showed that Nanog binds to the Tcf7 and interferes the interaction between β-catenin and Tcf7 *in vitro*, we then tested this possibility in *vivo*. We injected mRNAs encoding TCF7-ΔβBD (β-catenin binding domain deleted TCF7), TCF7-ΔGroBD (Groucho/Nanog binding domain deleted TCF7) and TCF7-ΔHMG (LEF1 binding domain deleted TCF7) into MZ*nanog* at one-cell stage. Theoretically, TCF7-ΔβBD can competitively bind to LEF1 without association of β-catenin, therefore resulting in decreased functional TCF7/β-catenin/LEF1 complex. Similarly, TCF7-ΔHMG can competitively bind with β-catenin but without the LEF1 association, therefore resulting in decreased level of functional TCF7/β-catenin/LEF1 complex. As shown in Figure 8a, overexpression of TCF-ΔβBD, TCF7-ΔHMG or Ctnnbip1 effectively rescued the hyper-dorsalized phenotype of MZ*nanog*, whereas overexpression of TCF7-ΔGroBD did not show any rescue (Fig. 8a). We then examined the *chd* expression of the recused MZ*nanog* embryos and found that overexpression of TCF7-βBD, TCF7-ΔHMG and Ctnnbip1 significantly rescued the hyper-dorsalization phenotype of MZ*nanog* which was characterized by a strong lateral and ventral extension of *chd* expression at the 30% epiboly stage (Fig. 8b). We further confirmed the rescue effect by TOPFlash luciferase assay (Fig. 8c).

**Figure 8.**
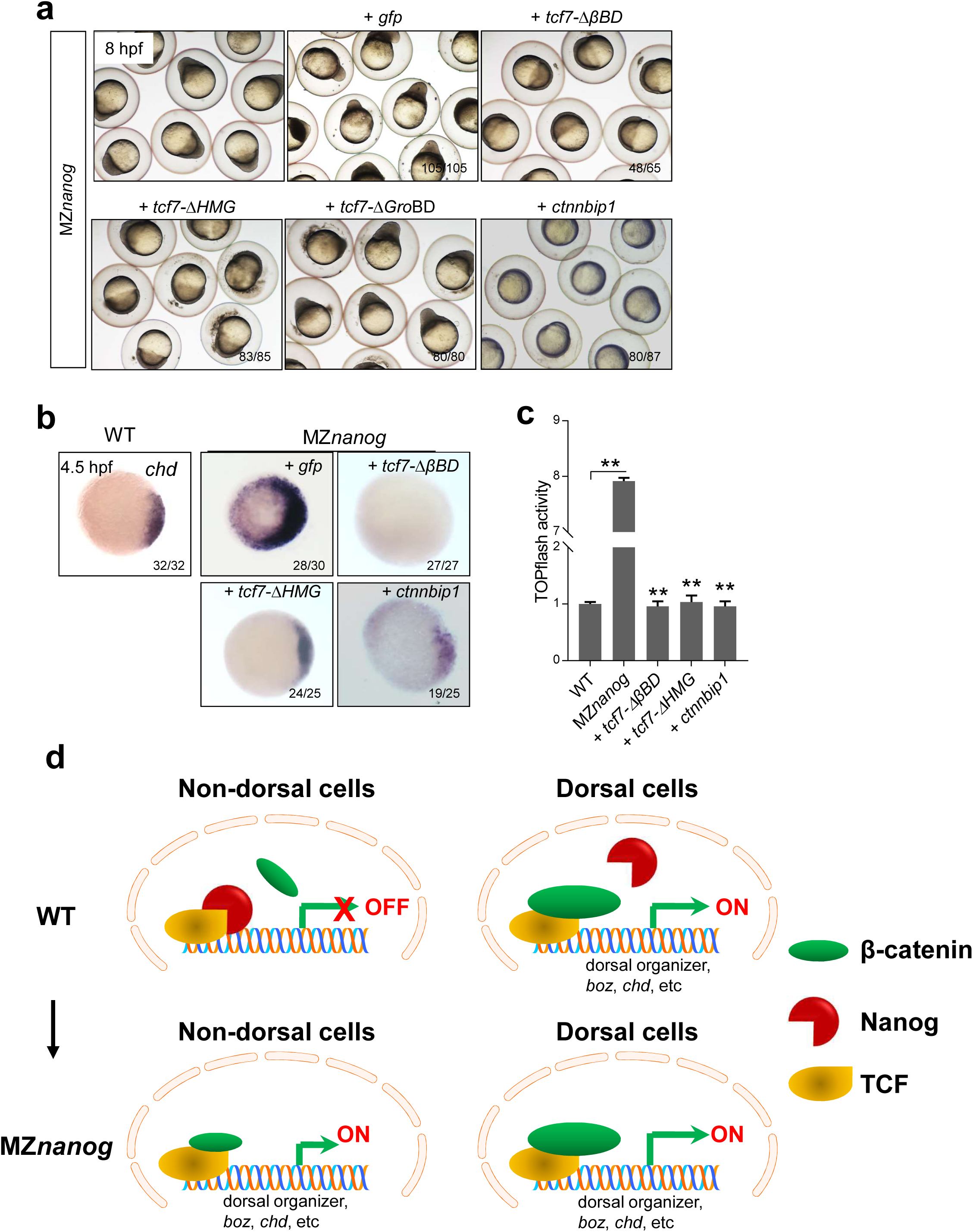
Confrontation of the β-catenin transcriptional activity in nucleus rescues the developmental defect of MZ*nanog*. (**a**) Overexpression of *tcf7-Δ*βBD, *tcf7-ΔHMG*, or *ctnnbip1* mRNA rescued the dorsalization phenotype of MZ*nanog*, while overexpression of *tcf7-ΔGroBD* did not. Phenotype was observed at 8 hpf. At least 50 embryos were injected and three independent experiments were performed. (**b**) Excessive and ectopic expression of *chd* in MZ*nanog* was rescued by overexpression of *tcf7-Δ*βBD, *tcf7-ΔHMG*, or *ctnnbip1*. WISH analysis of *chd* was detected at 4.5 hpf. (**c**) Detection of TOPflash activity in rescued MZ*nanog* embryos. ** means p < 0.01. (**d**) The model of Nanog repressing β-catenin transcriptional activity in non-dorsal cell nuclei in wildtype embryo and the ectopic activation of β-catenin transcriptional activity in the absence of Nanog.

All these results demonstrate that, in WT embryo, Nanog and β-catenin competitively combine with TCF7 and maintain the Wnt/β-catenin activity at homeostatic levels in different territories of the embryo. When Nanog is absent in MZ*nanog*, low amount of nuclear β-catenin in the non-dorsal-most cells can form functional β-catenin/TCF complexes and hyperactive the target genes, which resulted in hyper-dorsalization (Fig. 8d).

## Discussion

The induction of vertebrate dorsal axis relies on nuclear accumulation of maternally provided β-catenin in early embryo, which activates the dorsal genes, such as *boz*, *chd*, *sqt*, etc. in a cluster of dorsal-most cells [15]. These cells will develop into dorsal precursors, starting to establish embryonic dorso-ventral axis. The significance of maternal β-catenin in establishing early dorsal axis is supported by gain- and loss-of-function studies in zebrafish, e.g., overexpression of β-catenin causes dorsalization and β-catenin-related mutant *ichabod* shows serve ventralization [64, 65]. In the present study, we have detected low amount of nuclear β-catenin accumulation in the non-dorsal cells in zebrafish early embryo, which is consistent with the previous finding in *Xenpous* [48, 49]. Therefore, the nuclear accumulated β-catenin in non-dorsal cells must be properly repressed, in order to prevent the embryo from ectopic-activation of maternal β-catenin activity and hyper-dorsalization. Through a series of genetic and biochemical studies, we uncovered a novel repressor of maternal β-catenin activity, Nanog, which suppresses the transcriptional activity of Wnt/β-catenin signaling by competitive binding to transcription-activator type TCF7 and disrupting the functional β-catenin/TCF complex. Our study therefore establishes the maternal Nanog as a key factor that safeguards the embryo against global activation of maternal β-catenin activity by interfering with TCF factors.

In previous knockdown study, maternal Nanog has shown to play a key role in controlling the MZT, mainly including ZGA and the clearance of maternal mRNA, together with Pou5f1 and SoxB1 [40]. In our study, we have successfully generated the maternal zygotic mutants of *nanog*, MZ*nanog*, to uncover the developmental and molecular functions of Nanog. We have shown that the transcriptional activation of a series of zygotic genes, such as *blf, mir-430* and *mxtx2*, is defective in MZ*nanog*, and the clearance of maternal transcripts such as *sod1* and miR-430 sensitive reporter mRNA, are disrupted in MZ*nanog*. All these strongly support the previous finding that Nanog plays a critical role in mediating MZT. Recently, two independent studies by generating MZ*nanog* mutants report that maternal Nanog is also critical for extraembryonic tissue, embryo architecture and cell viability [66, 67]. In our study, we also observed those defects in MZ*nanog*, and more importantly we further reveal that all these defects could be perfectly rescued by overexpression of full-length Nanog, C-terminal truncated Nanog or a transcription-activator type Nanog (Vp16-Nanog). Therefore, our study demonstrates that Nanog acts as a transcriptional activator in mediating MZT, embryo architecture and cell viability.

Previous *in vitro* studies have shown that Groucho/TLE is a major co-repressor that can replace β-catenin from the TCF/Lef complex in the absence of Wnt ligands, resulting in transcriptional repression of Wnt/β-catenin signaling pathway [8, 10]. However, *in vivo* studies either by knockdown or knockout of zebrafish *tle* genes, no obvious early defects were detected, suggesting that Tles do not suppress maternal Wnt/β-catenin signaling during zebrafish early embryonic development. Although maternal Nanog is required for ZGA, to our surprise, the maternal Wnt/β-catenin targets, such as *boz* and *chd*, are not transcriptionally silenced or downregulated in MZ*nanog*. Instead, they are hyper-activated in the absence of maternal Nanog. Therefore, although those maternal Wnt/β-catenin targets are zygotic genes, they do not require the transcriptional activation by Nanog, and are mainly triggered by the maternal β-catenin transcriptional activity, which is repressed by maternal Nanog. Hence, the ubiquitous and high-level expression of Nanog in early embryo is not only required for MZT but also for the proper formation of dorso-ventral axis.

As a homeobox protein, zebrafish Nanog functions as a transcriptional activator, just like Oct4 and Sox2 in human [68]. In zebrafish MZ*nanog* embryo, although the Vp16-Nanog (N-terminal of Nanog replaced by strong transcriptional activator Vp16) could perfectly substitute endogenous Nanog in the aspects of controlling MZT, it did not suppress the elevated Wnt/β-catenin signaling activity in MZ*nanog* embryos. This indicates that the repression effect on Wnt/β-catenin signaling by Nanog is dependent on its N-terminal but independent of its transcriptional activation activity. Our study reveals that the N-terminal of Nanog could physically interact with the classical Groucho/TLE-binding domain of Tcf7, which interaction interferes the formation of TCF/β-catenin transcriptional activation complex. This conclusion is supported by rescuing the dorsalization phenotype of MZ*nanog* by overexpression of Tcf7-ΔβBD, Tcf7-ΔHMG or Ctnnbip1, which can disrupt the functional β-catenin/TCF complex. Therefore, our study has uncovered a novel role of Nanog in embryonic development, that Nanog controls the maternal Wnt/β-catenin transcriptional activity inside the nucleus to safeguard the embryo against global activation of maternal Wnt/β-catenin and to form the primary dorso-ventral axis. In view of the key role of Nanog in activating the first wave of zygotic genes and clearance of maternal mRNA [40], our study further establishes the central role of Nanog in coordinating early embryonic development of vertebrates.

## Acknowledgement

We thank Kuoyu Li from the China Zebrafish Resource Center (CZRC) for zebrafish raising and management. We thank Fang Zhou from Analysis and Testing Center of Institute of Hydrobiology, CAS for technique support of confocal imaging, Dr. Yun Liu from Sun Yat-sen University for denoting miR430 mimics and GFP-3xIPT-miR-430 reporter, Prof. Anming Meng from Tsinghua University for providing pCS2+*hwa* construct. This work was supported by the National Natural Science Foundation of China (grant Nos: 31721005, 31671501 and 31702323), the National key R&D Program of China (2018YFA0801000), the Youth Innovation Promotion Association of Chinese Academy of Sciences and the State Key Laboratory of Freshwater Ecology and Biotechnology (grant number 2019FBZ05).

## Author contributions

M.H. conceived the study, performed experiments, analyzed data and drafted the paper. R.Z., F.Z. and S.J. performed experiments. H.W. contributed essential reagents. D.Y. analyzed data. Y.S. designed the project, analyzed and interpreted the data, revised and finalized the manuscript.

## METHODS

### Zebrafish maintenance

All the zebrafish used in this study were maintained and raised as previously described [69] at the China Zebrafish Resource Center (Institute of Hydrobiology, CAS, Wuhan, China). The wild-type embryos were collected by natural spawning from AB strain. The experiments of using zebrafish were performed under the approval of the Institutional Animal Care and Use Committee of the Institute of Hydrobiology, CAS.

### Generation of *nanog* mutants

*nanog* mutants were generated by TALENs as previously described [54]. A pair of PCR primers, nanog-WT-F1: 5′-ACCCATCTTATCATGCATAT-3′ and nanog-TALEN-R2: 5′-TATCGCGTCGAGTGTACGCATG-3′, was used to screen and distinguish *nanog* homozygous and heterozygous. The maternal-zygotic *nanog* mutant line (MZ*nanog*) was derived and kept by injection of *nanog* mRNA at one-cell stage or derived from crossing of *nanog* female heterozygous with *nanog* male homozygous.

### Morpholino sequences and injections

Morpholinos were obtained from Gene Tools, LLC. The sequence of the morpholinos used is: *nanog* MO: 5′-CTGGCATCTTCCAGTCCGC<u>CAT</u>TTC-3′ (translation-blocking MO covering the translation initiation start, underlined).

*wnt8a* MO1: 5′-ACGCAAAAATCTGGCAAGGGTTCAT-3′ [70], *wnt8a* MO2: 5′-GCCCAACGGAAGAAGTAAGCCATTA-3′ [70], *tcf7l1a* MO: 5′-CTCCGTTTAACTGAGGCATGTTGGC-3′ [53], *tle3b* (*gro1*) MO: 5′-CGGCCCTGCGGATACATCTTGAATG-3′ [71], and *tle3a* (*gro2*) MO: 5′-ATGTATCCTTTATTTATTGGAGCTC-3′ [72] have been described previously.

For all experiments, MO or combinations of MOs were injected in more than 50 embryos and experiments were reproduced at least three times.

Amount of MO injected in this study: *nanog* MO (low dose): 0.5 ng/embryos, *nanog* MO (moderate dose): 1.2 ng/embryo, *nanog* MO (no phenotype): 160 pg/embryo, *wnt8a*1 MO: 500 pg/ embryo, *wnt8a2* MO: 500 pg/ embryo, *tcf7l1a* MO (head truncated): 1.6 ng/ embryo, *tcf7l1a* MO (no phenotype): 800 pg/embryo, *gro1* MO: 1.8 ng/embryo, *gro2* MO: 1.8 ng/embryo. 1 μL sample was injected into approximate 1000 embryos.

### mRNA synthesis and injection

For capped mRNA synthesis, full-length, mutated or truncated cDNAs have been cloned into pCS2+ vector, subsequently linearized with *Not*I and transcribed using SP6 RNA polymerase using the mMESSAGE mMACHINE kit from Ambion and injected into one-cell stage embryos if not specified. All injection experiments were performed on more than 50 embryos and reproduced at least three times.

Amount of mRNA injected in this study: *wnt8a* mRNA (head truncated): 1 ng/μL (1 pg/embryo), *wnt8a* mRNA (no phenotype): 0.1 ng/μL, β-catenin2 mRNA: 200 ng/μL, *nanog_*FL mRNA: 500 ng/μL, *nanog*_ΔCT mRNA: 25 ng/μL, *vp16*-*nanog* mRNA: 250 ng/μL, *mxtx2* mRNA: 1 ng/μL, *hwa* mRNA: 200 ng/μL, *tcf7-ΔβBD* mRNA: 800 ng/μL, *tcf7-ΔHMG* mRNA: 800 ng/μL, *tcf7l1a* mRNA: 400 ng/μL, *gsk3b* mRNA: 900 ng/μL, *ctnnbip1* mRNA: 400 ng/μL, *cby1* mRNA: 200ng/uL. GFP mRNA (100 ng/μL) was injected as control for rescue experiments.

For localized injection, dechorioned embryos were collected at 16-32 cell stage and injected twice, one is at margin cell, another is at the middle cell of blastula. β-catenin mRNA or *hwa* mRNA were co-injected with *mCherry* mRNA, and *mCherry* was used as injection indicator.

For miR-430 mimics injection, a mixture of miR-430a, miR-430b and miR-430c was injected at 3.3 μmol/μL. The miR-430 mimics and GFP-3xIPT-miR-430 reporter was a gift by Dr. Yun Liu and injected as described [73]. 1 μL sample was injected into approximate 1000 embryos.

### *In situ* hybridization

PCR-amplified sequences of genes of interest were used as templates for the synthesis of an antisense RNA probe, labelled with digoxigenin-linked nucleotides. whole-mount *In situ* hybridization (WISH) on embryos were performed as described[74]. For *in situ* hybridization on frozen section, adult ovaries were stripped and fixed overnight with 4% PFA in PBS at 4°C, then embedded with OCT and dissected at 10um. The procedures of hybridization followed Wilkinson, et al [75].

### Immunofluorescence

Whole-mount immunofluorescence was carried out based on the standard protocol using the following primary antibodies: mouse anti–β-catenin antibody (C7267, sigma, 1:500 dilution), The rabbit anti-Nanog polyclonal antibody was customized by ABclone using a recombinant full-length of Nanog protein (1:200 dilution). For nuclear β-catenin staining, embryos were fixed overnight with 4% PFA in PBS at 4 °C, and washed with PBST (0.1% Triton X-100 added) and permeabilized with distilled water for one hour. The FITC conjugated goat anti-mouse or rabbit antibody (Thermo, 1:500) was used as secondary antibody. After four times of washing, using PBST with 0.1% Triton X-100, embryos were incubated in DAPI solution (5 ug/mL in PBST) for 1 hour at room temperature. Then, embryos were washed and mounted for observation. Immunostained embryos were photographed using a Leica SP8 confocal microscope.

### Luciferase assay

15 pg of Topflash construct and 1.5 pg of Renilla reporter were mixed and co-injected into 1-cell stage embryos. 4 ng of Topflash construct and 0.4 ng of Renilla reporter were mixed and co-transfected into 293T cells with indicated plasmids. Injected embryos were collected at 8 hours post fertilization (hpf) and transfected cells were collected at 24 hours after transfection. The luciferase activity was measured by Dual-Luciferase Reporter Assay System (Promega, USA). TopFlash assays were performed in triplicate for each sample. Student’s t-test was used to assess the statistical significance.

### Stem-loop RT-PCR

Stem-loop RT-PCR was performed as previously described [76] to quantify the expression of miR-430. Total RNAs were reversely transcribed using the miR-430-RT primer 5’-GTCGTATCCAGTGCAGGGTCCGAGGTATTCGCACTGGATACGACCTACCCCA-3’ and the U6 RT primer 5’-AAAAATATGGAGCGCTTCACG-3’. The PCR primers were listed as follows: for U6, forward primer 5′-TTGGTCTGATCTGGCACATATAC-3′ and reverse primer 5′-AAAAATATGGAGCGCTTCACG-3′; for *miR-430a*, forward primer 5′-GCGAAGTGCTATTTGTTGGGGT-3′ and reverse primer 5′-GTGCAGGGTCCGAGGT-3′, for *miR-430b*, forward primer 5′-GCGTGCTATCAAGTTGGGGTAG-3′ and reverse primer 5′-GTGCAGGGTCCGAGGT-3′.

For normal RT-PCR, Total RNAs were reversely transcribed using oligo dT, and PCR primers were listed as follows: *tle3a*: forward primer 5′-TGGCCTCGTCTGGCAGTA-3′ and reverse primer 5′-CGAAGGGAGTTGGGTAAGTG-3′, *tle3b*: forward primer 5′-GGACCTTCATAACCAAACCC-3′ and reverse primer 5′-TTCCCACAGCCAGCCACT-3′, *boz*: forward primer 5′-CGTTCCAGTCTCCTACTAC-3′ and reverse primer 5′-GCAGGTTGTCTGTCTCTT-3′, *wnt8a1*: forward primer 5′-GCGTCGTTGGTTATGTCT-3′ and reverse primer 5′-AACTCCAGCAGCACTTATAG-3′, *wnt8a2*: forward primer 5′-GGAGGATGTAGCGACAAC and reverse primer 5′-TTGCCAATCTCACGGAAG-3′. Real-time PCR was performed using the SYBRGreen Supermix from BioRad (USA) on a BioRad CFX96.

### Immunoprecipitation and Western blot

For immunoprecipitation assays, the different full length and truncated cDNAs have been cloned into pCMV-myc (N-terminal myc tag) or in pCGN-HAM (N-terminal multiple HA tag) vectors. HEK293T cells were transiently transfected with the indicated constructs of interest using VigoFect (Vigorous Biotechnology, China). 24 hours after transfection, cells were harvested and lysed in RIPA buffer (50mM Tris-HCl [pH 7.4], 150mM NaCl, 1% NP40, 0.25% C24H39O4Na, 1mM EDTA, 1mM NaF, and Protease Inhibitor Cocktail [Sigma]). Coimmunoprecipitation experiments were performed as described previously[77].

Primary antibodies and dilutions for western blot were: Myc (Santa Cruz Biotechnology, 1:2000), HA (Sigma, 1:5000), Nanog (ABclone, 1:2000), total β-catenin (Sigma, 1:5000), active β-catenin (CST, 1:1000), Signals were detected with ECL Western blotting detection reagents (Millipore, Germany).

### Statistical analysis

Significance of differences between means was analyzed using Student’s t *test*. P value below 0.05 marked as *, and P value below 0.01 marked as **, NS means no significant difference.

## Supplementary Information

**S1 Fig.**
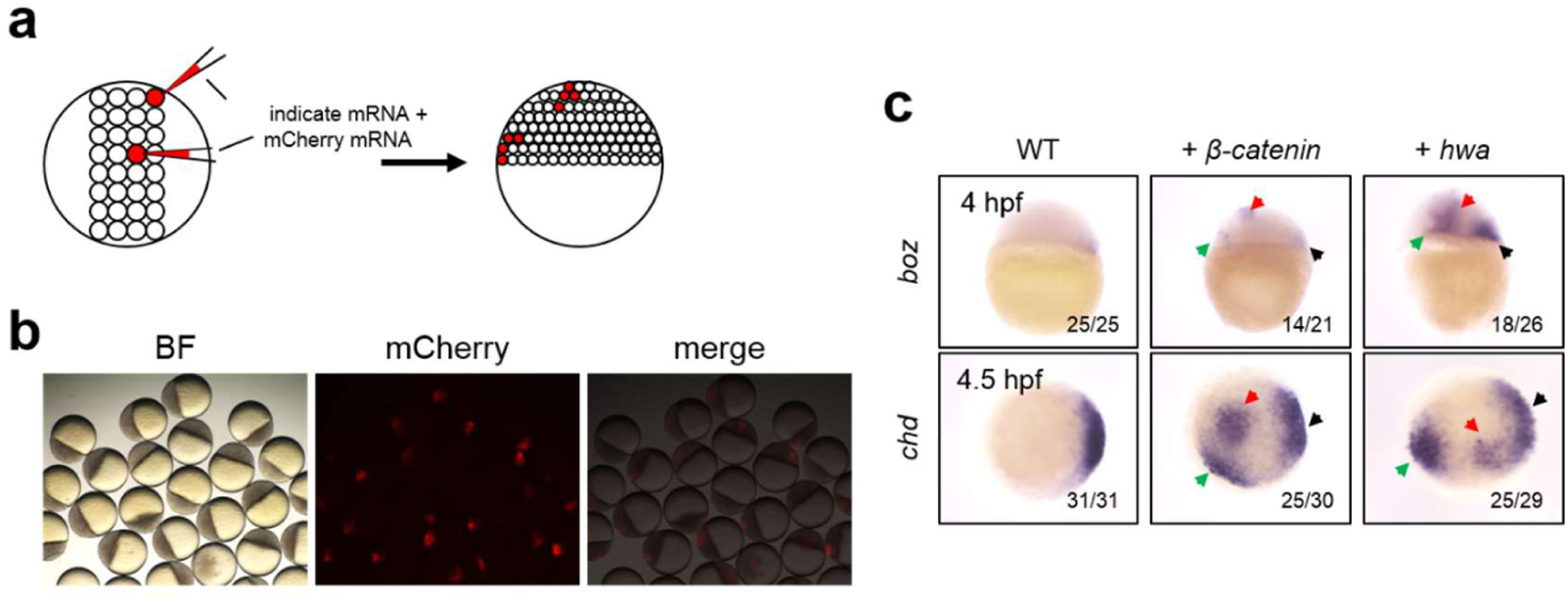
Localized injection of *β-catenin* or *hwa* mRNA induce ectopic dorsal organizer. **(a)** A diagram of localized injection into two cells at 32-cell stage. One injected cell is on the margin, and the other injected cell is located at the center of blastula. **(b)** *β-catenin* or *hwa* mRNA was co-injected with mCherry mRNA and located injected mCherry fluorescence were observed at 4hpf. **(c)** Detection of ectopic expression of *boz* and *chd* induced by *β-catenin* and *hwa* mRNA. Embryos with fluorescence were collected and examined by WISH. *boz* was detected at 4hpf, *chd* was detected at 4.5hpf.

**S2 Fig.**
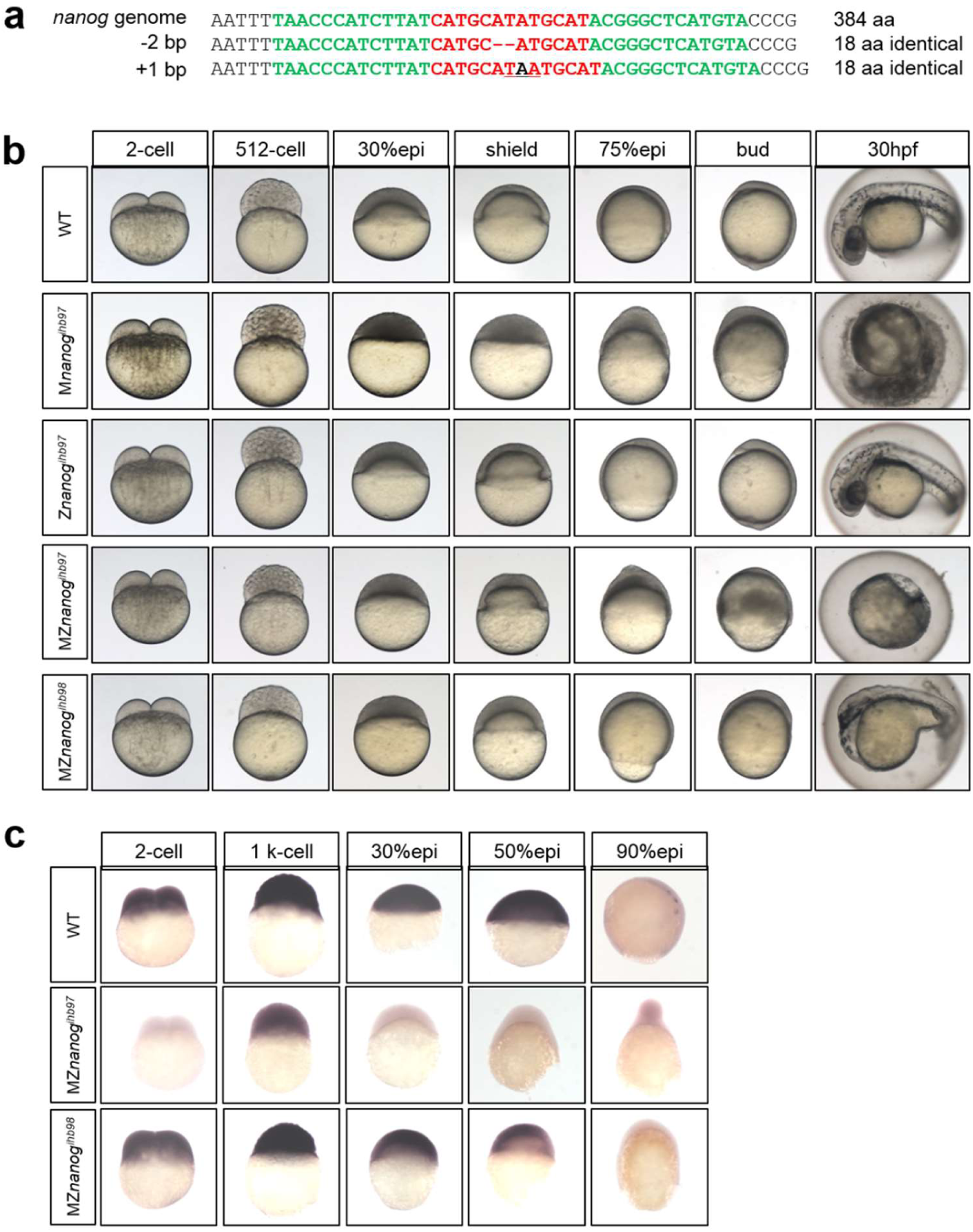
Characterization of *nanog* mutants. **(a)** Generation of two *nanog* mutant alleles by TALEN. TALEN left and right arm is marked in green and target sequence is in red. Two different mutant lines were obtained, 2-bp deletion line (*nanog^ihb97^*) and 1-bp insertion line (*nanog^ihb98^*). 2-bp deletion leads to frame-shift of Nanog protein, and 1-bp insertion results in premature termination of Nanog protein at the mutation site. (b) Phenotype characterization of M*nanog^ihb97^*, Z*nanog^ihb97^*, and two different mutants, MZ*nanog^ihb97^* and MZ*nanog^ihb98^*. Both of M*nanog^ihb97^* and two types of maternal and zygotic *nanog* mutants, MZ*nanog^ihb97^*, MZ*nanog^ihb98^* show slowly development and abnormal cell movement, then die within 24hpf. however, Z*nanog^ihb97^* mutant shows normal development and reproduction, as the same as WT. (c) Maternal *nanog* expression disappeared and small amount of zygotic *nanog* was detected in MZ*nanog^ihb97^*. Both low expression of maternal and zygotic *nanog* were detected in MZ*nanog^ihb98^*.

**S3 Fig.**
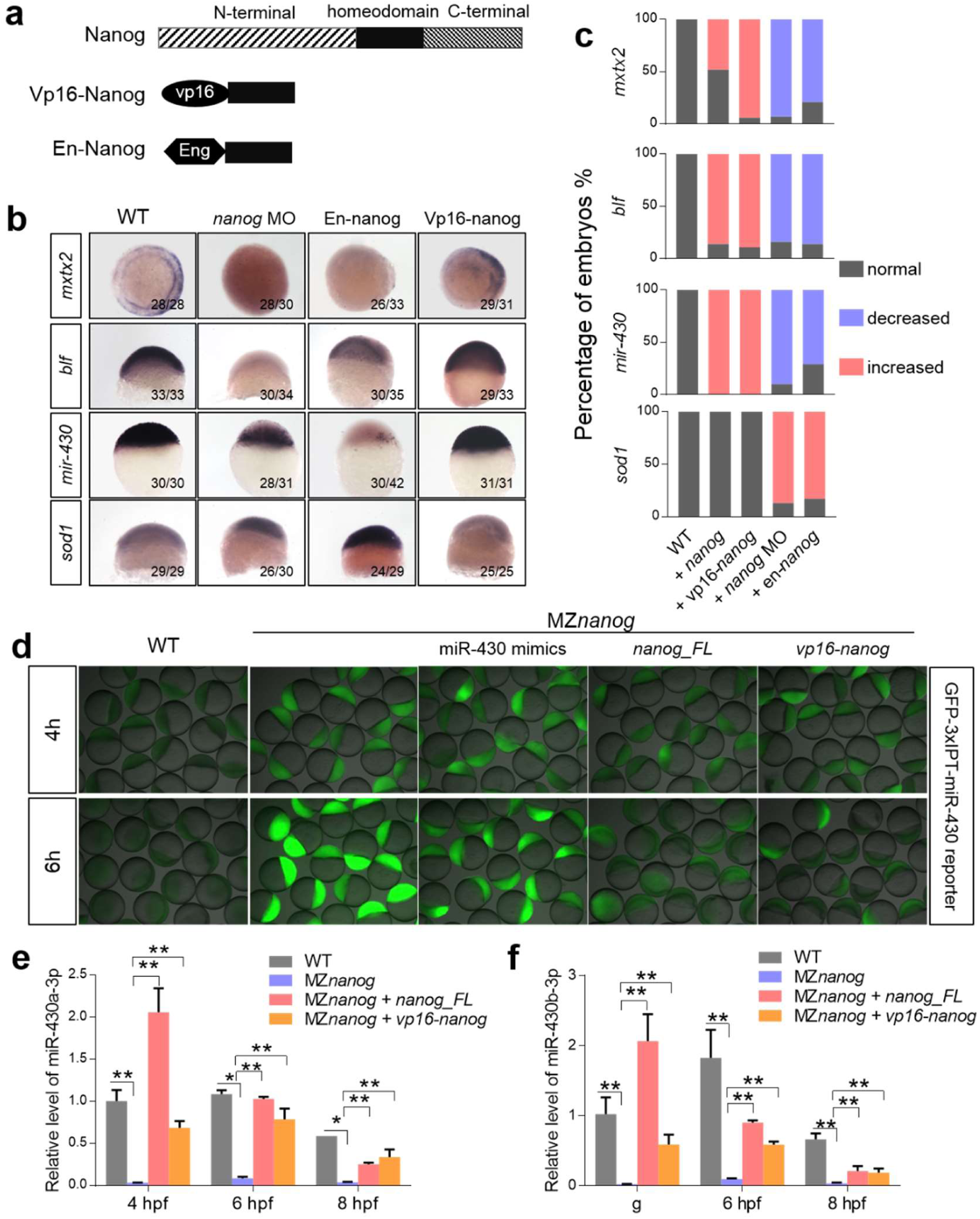
Vp16-Nanog could fully rescue the MZT defects in MZ*nanog*. **(a)** The diagram of Vp16-Nanog (vp16-Nanog homeodomain) and En-Nanog (Enrgrailed2-Nanog homeodomain). The transcription activator VP16 or repressor Enrgrailed2 was fusion with Nanog homeodomain to result in Vp16-Nanog or En-Nanog. (b) The expression of mesendoderm marker, *mxtx2* (which is also a direct target of Nanog), zygotic gene, *blf*, and micro-430 precursor, *mir-430*, showed decreased or absent in *nanog* morphants and En-Nanog injected embryos, but enhanced in *vp16-nanog* overexpressed embryos. The expression of miR-430 target gene, *sod1*, failed to be cleared in *nanog* morphants and En-Nanog injected embryos at shield stage. (c) The percentage of embryos counted in WISH of (b). (d) The fluoresce intensity of GFP-3xIPT-miR-430 reporter which carries a target sequence of miR-430 is negative correlated with the expression of miR-430. The intensity of GFP was higher in MZ*nanog* than WT, indicating deletion of *nanog* resulted in inactivation of miR-430 expression. Meanwhile, overexpression of miR-430 mimics, *nanog* or *vp16-nanog* in MZ*nanog* restored the high expression of GFP-3xIPT-miR-430 reporter. (e) and (f) miR-430a and miR-430b are inactivated in MZ*nanog*, overexpression of *nanog_*FL or *vp16-nanog* can fully restore the expression failure of miR-430a and miR-430b at 4, 6, 8 hpf. * means p<0.05, ** means p < 0.01.

**S4 Fig.**
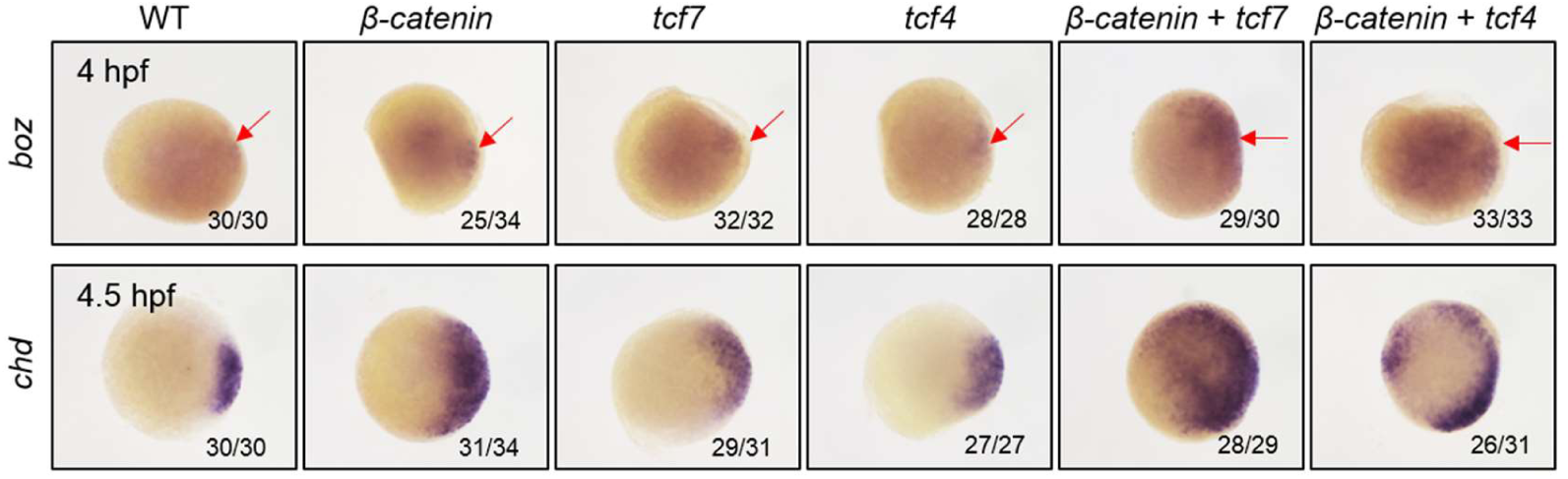
Tcf7 and Tcf4 are activator-type TCFs in zebrafish. Injection of low dose of β-catenin mRNA (200 pg/embryo), *tcf7* mRNA, or *tcf4* mRNA alone, or co-injection of low dose of *β-catenin* mRNA (200 pg/embryo) with *tcf7*, or *tcf4*, at 1-cell stage and detection the expression of *boz* and *chd*. Both individual injection and co-injection of TCF with β-catenin induced up-regulation of *boz* and *chd*, indicating that *tcf7* and *tcf4* acts as activator-type TCF in Wnt/β-catenin pathway.

**S5 Fig.**
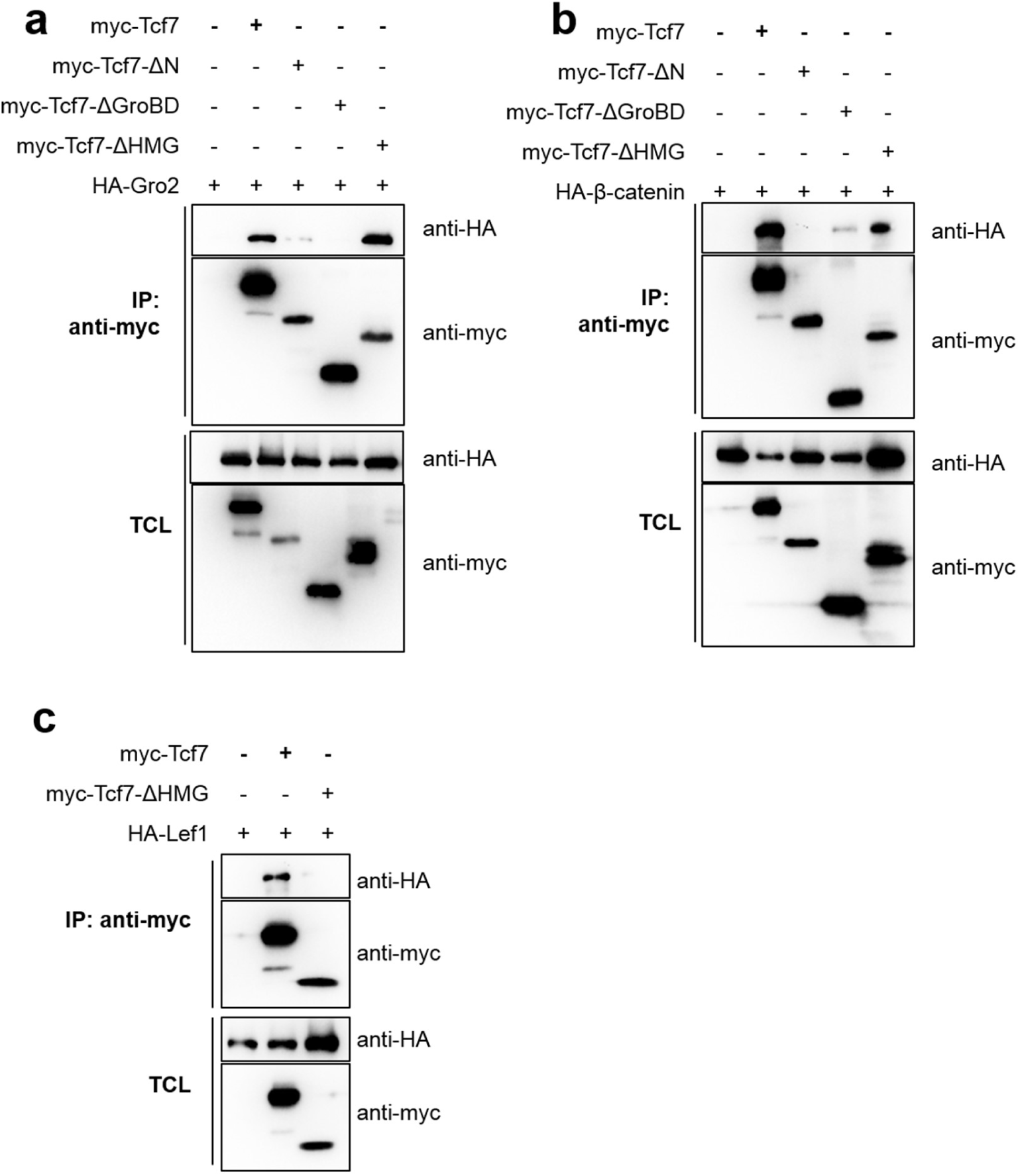
Interaction of TCF7 with Groucho/TLE, β-catenin and LEF1. **(a)** TCF7 interacts with Groucho/TLE through predicted Groucho/TLE-binding domain. Different Myc-tagged Tcf7 were constructed and co-transfection with HA-Groucho2/TLE3a in HEK293T cells. Deletion of Tcf7 predicted Groucho/TLE-binding domain disrupts its interaction with Groucho2/TLE3a, indicating that Groucho2/TLE3a physically binds with the potential Groucho/TLE-binding domain of Tcf7. (b) TCF7 interacts with β-catenin through its N-terminal. Deletion of Tcf7 N-terminal disrupts its interaction with β-catenin, indicating that β-catenin physically interacts with the N-terminal of Tcf7. (c) TCF7 interacts with LEF1 through its HMG domain (C-terminal).

